# Differential Requirements for the RAD51 Paralogs in Genome Repair and Maintenance in Human Cells

**DOI:** 10.1101/609115

**Authors:** Edwige B. Garcin, Stéphanie Gon, Rohit Prakash, Meghan R. Sullivan, Gregory J. Brunette, Anne De Cian, Jean-Paul Concordet, Carine Giovannangeli, Wilhelm G. Dirks, Sonja Eberth, Kara A. Bernstein, Maria Jasin, Mauro Modesti

## Abstract

Deficiency in several of the classical human RAD51 paralogs [RAD51B, RAD51C, RAD51D, XRCC2 and XRCC3] is associated with cancer predisposition and Fanconi anemia. To investigate their functions, isogenic disruption mutants for each were generated in non-transformed MCF10A mammary epithelial cells and in transformed U2OS and HEK293 cells. In U2OS and HEK293 cells, viable ablated clones were readily isolated for each RAD51 paralog; in contrast, with the exception of RAD51B, RAD51 paralogs are cell-essential in MCF10A cells. Underlining their importance for genomic stability, mutant cell lines display variable growth defects, impaired sister chromatid recombination, reduced levels of stable RAD51 nuclear foci, and hypersensitivity to mitomycin C and olaparib. Altogether these observations underscore the contributions of RAD51 paralogs in diverse DNA repair processes, and demonstrate essential differences in different cell types. Finally, this study will provide useful reagents to analyze patient-derived mutations and to investigate mechanisms of chemotherapeutic resistance deployed by cancers.

## INTRODUCTION

RecA family recombinases are universally found in the three domains of life: RadA in Archaea, RecA in Bacteria and RAD51/DMC1 in Eukarya (Lin et al. 2006). By coordinating ATP, they assemble nucleoprotein filaments on single-stranded DNA that promote DNA sequence homology recognition in duplex DNA and catalyze strand exchange (Bianco et al. 1998). These enzymes play a central role in the maintenance of genome integrity as the DNA transactions they support are essential for repair of DNA double-strand breaks (DSBs) by homologous recombination (HR), and in protection and rescue of stalled or collapsed DNA replication forks (Kolinjivadi et al. 2017). Interestingly, additional RecA-like genes have evolved in the three domains of life presumably after duplication and divergent evolution of a common ancestor (Lin et al. 2006; Thacker 1999). While structurally related, these paralogs do not promote homology recognition and strand exchange but rather act in part as accessory factors to the core recombinases.

In human cells and vertebrates in general, besides RAD51 and DMC1, six RAD51 paralogs have been identified. RAD51B, RAD51C and RAD51D were discovered based on DNA sequence alignments, and XRCC2 and XRCC3 through functional complementation of the ionizing radiation (IR) sensitivity of Chinese hamster mutant cells (Albala et al. 1997; Cartwright et al. 1998; Dosanjh 1998; Pittman et al. 1998; Tebbs et al. 1995). These five RAD51 paralogs (herein referred to as the classical RAD51 paralogs and focus of this study) are believed to form two functionally distinct heterotypic complexes: the RAD51B-RAD51C-RAD51D-XRCC2 complex with sub-complexes RAD51B-RAD51C and RAD51D-XRCC2; and the RAD51C-XRCC3 complex (Figure 1A) (Braybrooke et al. 2000; Liu 2002; Masson et al. 2001b, 2001a; Miller et al. 2002; Schild et al. 2000; Sigurdsson et al. 2001; Wiese et al. 2006; Yokoyama et al. 2004; Yonetani et al. 2005). Lastly, SWSAP1, a non-classical RAD51 paralog, was recently identified as a component of the Shu complex (Liu et al. 2011b; Martino and Bernstein 2016).

**Figure 1.**
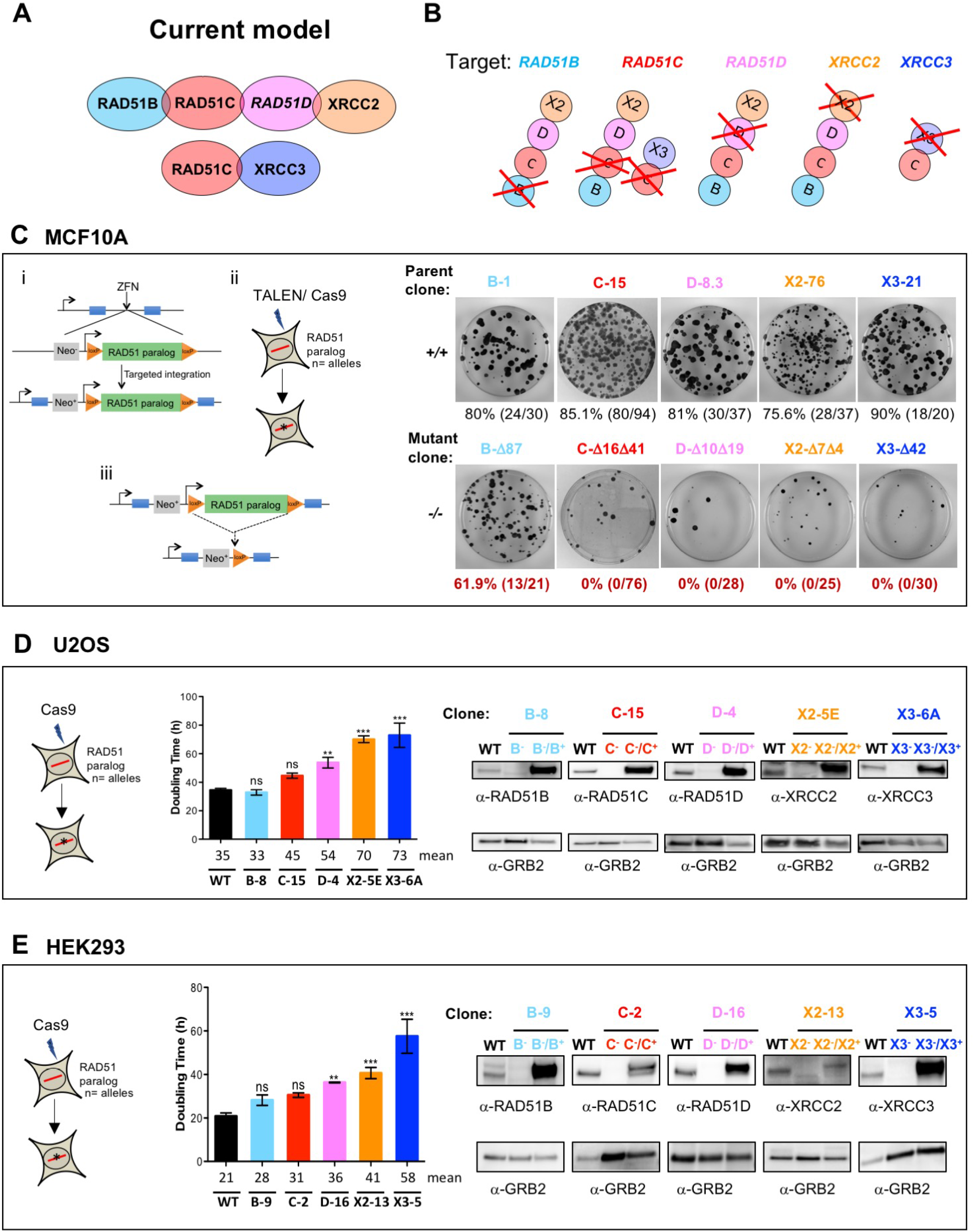
Individual disruption of the classical RAD51 paralogs in human MCF10A, U2OS, and HEK293 cells reveal variable effects on cell viability. **(A)** According to the current model based on biochemical studies, RAD51 paralogs exist in two different complexes where RAD51C is a common member of both. **(B)** Schematics showing the individual RAD51 paralog knockout. **(C) i** RAD51 paralog cDNA flanked by loxP sites was first targeted into the AAVS1 locus using zinc finger nucleases in MCF10A cells and the correct integration was selected as it results in expression of the promoterless Neo gene (these cells are designated as parental conditional). **ii** Nuclease-mediated gene editing was used on these parental conditional cells to mutate the endogenous RAD51 paralog alleles. **iii** Isogenic parental and conditional mutant RAD51 paralog cells were infected with lenti-Cre to delete the RAD51 paralog cDNA, post two days Cre infection cells were diluted for colony formation and analysis. Right panels shows examples of the Giemsa stained plates from the various isogenic parental (+/+) and knockout (-/-) cell lines after Cre treatment and underneath shows the number and % Cre excised clones for each. The conditional parental and mutant RAD51 paralog clones that were used in this study are listed on top of each plate. **(D, E)** CRISPR/Cas9 mediated gene editing was used on U2OS and HEK293 cells to delete the RAD51 paralog endogenous alleles. Graphs show doubling time for the wild-type and the mutant RAD51 paralog cells. Number of viable cells was estimated using the CellTiter-Glo Luminescent Cell Viability Assay at 24 h, 48 h, 72 h and 96 h after seeding. Cell doubling time was calculated by nonlinear regression (exponential growth equation) analysis. Results are presented as means +/− SD from three experiments. Comparisons between mutant and wild-type cells were analyzed by performing unpaired one-way ANOVA followed by Tukey’s test. ^ns^ p > 0.05, ** p ≤ 0.01, *** p ::: 0.001. Immunoblots of crude cellular extracts from wild-type cells, mutant cells and mutant cells stably complemented with a retroviral construct expressing the corresponding wild-type allele are shown on the right side. GRB2 was used as loading control. The mutant RAD51 paralog clones for U2OS and HEK293 that were used in this study are listed on top of each immunoblot.

To begin to decipher the specific functions of the five classical RAD51 paralogs, all but XRCC3 have been ablated in mice (Deans et al. 2000; Pittman and Schimenti 2000; Shu et al. 1999; Smeenk et al. 2010). In each case, ablation of RAD51 paralogs results in embryonic lethality or early neonatal death, precluding any detailed phenotypic analysis (Prakash et al. 2015). Nevertheless, previous cellular studies delineated critical roles for the classical RAD51 paralogs in genome maintenance. These cellular studies include studies using 1) Chinese hamster mutant cells deficient in RAD51C, RAD51D, XRCC2 or XRCC3; 2) DT40 chicken B-lymphocyte cells individually ablated for each of the five classical RAD51 paralogs; and 3) cells derived from the RAD51 paralog mutant mice (Adam et al. 2007; Bishop et al. 1998; French et al. 2002; Godthelp 2002; Hinz et al. 2006; Johnson et al. 1999; Liu et al. 1998; Pierce et al. 1999; Takata et al. 2001, 2000; Tambini et al. 1997; Tebbs et al. 1995). Typically, classical RAD51 paralog deficient cells are sensitive to DNA damaging agents including ionizing radiation (IR) and DNA crosslinking agents such as mitomycin C (Liu et al. 1998; Takata et al. 2001, 2000; Tebbs et al. 1995; Yonetani et al. 2005). Mutant cells display increased spontaneous chromosomal abnormalities, decreased frequencies of DNA damage-induced sister chromatid exchanges, reduced DNA damage-induced RAD51 focus formation, and deficiencies in replication fork protection (Badie et al. 2009; Bishop et al. 1998; Cui et al. 1999; Date et al. 2006; Deans et al. 2003; French et al. 2002; Griffin et al. 2000; Johnson et al. 1999; Katsura et al. 2009; Liu et al. 1998; Pierce et al. 1999; Rodrigue et al. 2006; Smiraldo et al. 2005; Somyajit et al. 2015; Sung et al. 2003; Takata et al. 2001; Yoshihara et al. 2004; Saxena et al. 2018). Overexpression of the core RAD51 recombinase partially suppresses DNA damage sensitivity of chicken classical RAD51 paralog mutant cells, suggesting a link with RAD51 function (Takata et al. 2001). Overall, these studies indicate that deficiencies in the classical RAD51 paralogs lead to genomic instability caused by compromised regulation of the core RAD51 recombinase.

Except for *RAD51B*, RAD51 paralog mutants have been isolated and characterized in Chinese hamster cells. In human cells, however, only *XRCC3* ablated cells generated in the human colon carcinoma HCT116 cell line have been reported to date (Yoshihara et al. 2004). Attempts to generate *RAD51B* or *RAD51C* mutants in HCT116 cells were not successful (Date et al. 2006; Katsura et al. 2009). As an alternative to study the function of the classical RAD51 paralogs in human cells, depletion approaches using small interfering RNA (siRNA) have been used (Chun et al. 2013; Jensen et al. 2013). These siRNA depletion experiments helped determine the genetic interactions between the RAD51B-RAD51C-RAD51D-XRCC2 and RAD51C-XRCC3 complexes, and with BRCA2, an important mediator of RAD51 function (Roy et al. 2011). Moreover, the classical RAD51 paralogs also have roles not directly linked to the control of RAD51 *per se.* Indeed, RAD51C is implicated in cell cycle checkpoint (Badie et al. 2009; Lio et al. 2004; Rodrigue et al. 2013), RAD51D and XRCC3 in telomere maintenance (Compton et al. 2007; Tarsounas and West 2005), and RAD51C, XRCC2 and XRCC3 in termination of gene conversion (Nagaraju et al. 2009, 2006; Puget et al. 2005).

Recently, mutations in the classical RAD51 paralog genes have been linked to predisposition to breast, ovarian or other cancers (Akbari et al. 2010; Golmard et al. 2013; Kondrashova et al. 2017; Loveday et al. 2011, 2012; Meindl et al. 2010; Orr et al. 2012; Osorio et al. 2012; Park et al. 2012; Somyajit et al. 2012; Zheng et al. 2010). Moreover, hypomorphic mutations in *RAD51C* and *XRCC2* confer Fanconi anemia disorder and are now named FANCO and FANCU, respectively (Park et al. 2016; Somyajit et al. 2010; Vaz et al. 2010). Obtaining human cell lines disrupted for the classical RAD51 paralog would therefore greatly help research aimed at understanding the impact of patient-derived mutations.

Despite more than two decades of research, our understanding of the specific functions of the five classical RAD51 paralogs and their complexes remains incomplete and controversial. To perform a comprehensive analysis of the roles of the classical RAD51 paralogs in human cells, we report here the generation and phenotypic characterization of individual RAD51 paralog conditional mutants in non-transformed MCF10A mammary epithelial cells, as well as viable human embryonic kidney HEK293 cells and U2OS osteosarcoma cells in which the five classical RAD51 paralogs have been individually disrupted (Figure 1B).

## RESULTS

### RAD51C, RAD51D, XRCC2 and XRCC3 are essential for survival in MCF10A but RAD51B is dispensable

To investigate the molecular functions of the classical RAD51 paralogs in a non-cancerous cell line with a near normal karyotype background, we chose the human mammary epithelial cell line MCF10A that contains an integrated DR-GFP HR reporter (Feng and Jasin 2017). Since the knockouts of the classical RAD51 paralogs are embryonic lethal in mice, we considered that biallelic mutations in RAD51 paralogs may lead to cell death in non-transformed cells. Therefore, a generalized conditional strategy was implemented to test for cell lethality. An expression cassette for each RAD51 paralog cDNA flanked by LoxP sites was first introduced at the safe harbor *AAVS1* locus (Brunet et al. 2009; Hockemeyer et al. 2009) to generate parental conditional cells for each RAD51 paralog (Figure 1C). To generate isogenic RAD51 paralog knockouts at the endogenous loci — *RAD51B^−/−^, RAD51C^−/−^, RAD51D^−/−^, XRCC2^−/−^* and *XRCC3^−/−^*, nuclease-guided gene disruption was used to target the coding region close to the start codon (Figure 1C and Figure S1). For *RAD51B*, CRISPR-Cas9 was used with paired gRNAs to generate a deletion within the exon. For *RAD51C, RAD51D, XRCC2* and *XRCC3,* TALENs were used in which one DNA binding site was in an exon and the other was in the adjacent intron; with this design, the RAD51 paralog cDNA would not be targeted for cleavage. In all cases, clones with disrupting biallelic mutations were obtained (Figure 1C, Figure S1, Table S1).

A self-deleting Cre recombinase was expressed in the biallelic conditional mutants as well as the parental conditional cells to delete the RAD51 paralog cDNA from the AAVS1 locus and clonogenic survival was determined (Figure 1C). Fewer colonies were obtained from the biallelic mutants compared to the parental cells. For *RAD51C, RAD51D, XRCC2* and *XRCC3*, genotyping revealed that none of the surviving colonies had undergone Cre-mediated excision of the cDNA flanked by LoxP sites, indicating that these genes are essential for cell survival. Surprisingly, *RAD51B* colonies were obtained that had excised the cDNA. Although excised colonies were obtained at a lower frequency from the mutant *RAD51B* cells compared with the parental cells, the cells could be propagated in culture. These results demonstrate that while *RAD51C, RAD51D, XRCC2* and *XRCC3* are essential for the survival of non-transformed MCF10A cells, *RAD51B* is not.

### Individual disruption of the five classical RAD51 paralogs in human U2OS and HEK293 cells are viable

Given that CHO cells deficient for RAD51C, RAD51D, XRCC2 or XRCC3, and human HCT116 cells deficient for XRCC3 are viable, we attempted to disrupt each classical RAD51 paralog individually in transformed human osteosarcoma U2OS and human embryonic kidney HEK293 cells. As with MCF10A cells, we utilized cell lines that harbor a stably integrated HR reporter: U2OS-SCR 18, (Puget et al. 2005) and HEK293 DR-GFP (Esashi et al. 2005), hereafter designated U2OS and HEK293. Each gene was targeted using CRISPR-Cas9 such that any truncated protein produced would lack the putative functional Walker B domain. For the initial screen, HEK293 and U2OS cells were transiently transfected with the Cas9-GFP and gRNA expression vectors, and single cells isolated by fluorescence activated cell sorting (FACS) based on GFP expression. (400 GFP-positive cells were individually seeded and about 5% of the clones expanded). The clones were screened by immunoblotting for loss of protein expression and genotyped. U2OS and HEK293 mutant clones harboring only frame-shifting indel mutations and no wild-type alleles were retained for further analysis (Figure 1D, Figure S2; Figure 1E, Figure S3; Table S1). Thus, viable cells individually disrupted for each of the five classical RAD51 paralogs were obtained in both U2OS and HEK293 cell lines.

### Disruption of classical RAD51 paralogs affects U2OS and HEK293 cell growth and fitness

Although all five RAD51 paralog disrupted cells are viable in both U2OS and HEK293, each displays defects in cell growth and fitness. Cell proliferation rates were assessed by measuring ATP levels, an indicator of metabolically active cells. Except for *RAD51B* and *RAD51C* disruption, RAD51 paralog mutant lines displayed significantly longer doubling time than wild-type cells (∼1.2 to 2-fold for U2OS and ∼1.6 to 3-fold for HEK293; Figure 1D, Figure 1E). Further, disruption of RAD51 paralogs caused a marked decrease in plating efficiency in both U2OS and HEK293 lines which was reversed by re-expressing the appropriate wild-type RAD51 paralog (Figure 2A). The levels of spontaneous apoptosis and cell death were also assessed using Annexin V and 7-Aminoactinomycin D labeling and FACS analysis. In proliferating cells, a modest accumulation of dead cells was observed (<10%) with little or no induction of apoptosis (Figure 2B). Altogether these results show that, while not essential for cell viability, classical RAD51 paralog disruption affects the basal growth properties of U2OS and HEK293 cells.

**Figure 2.**
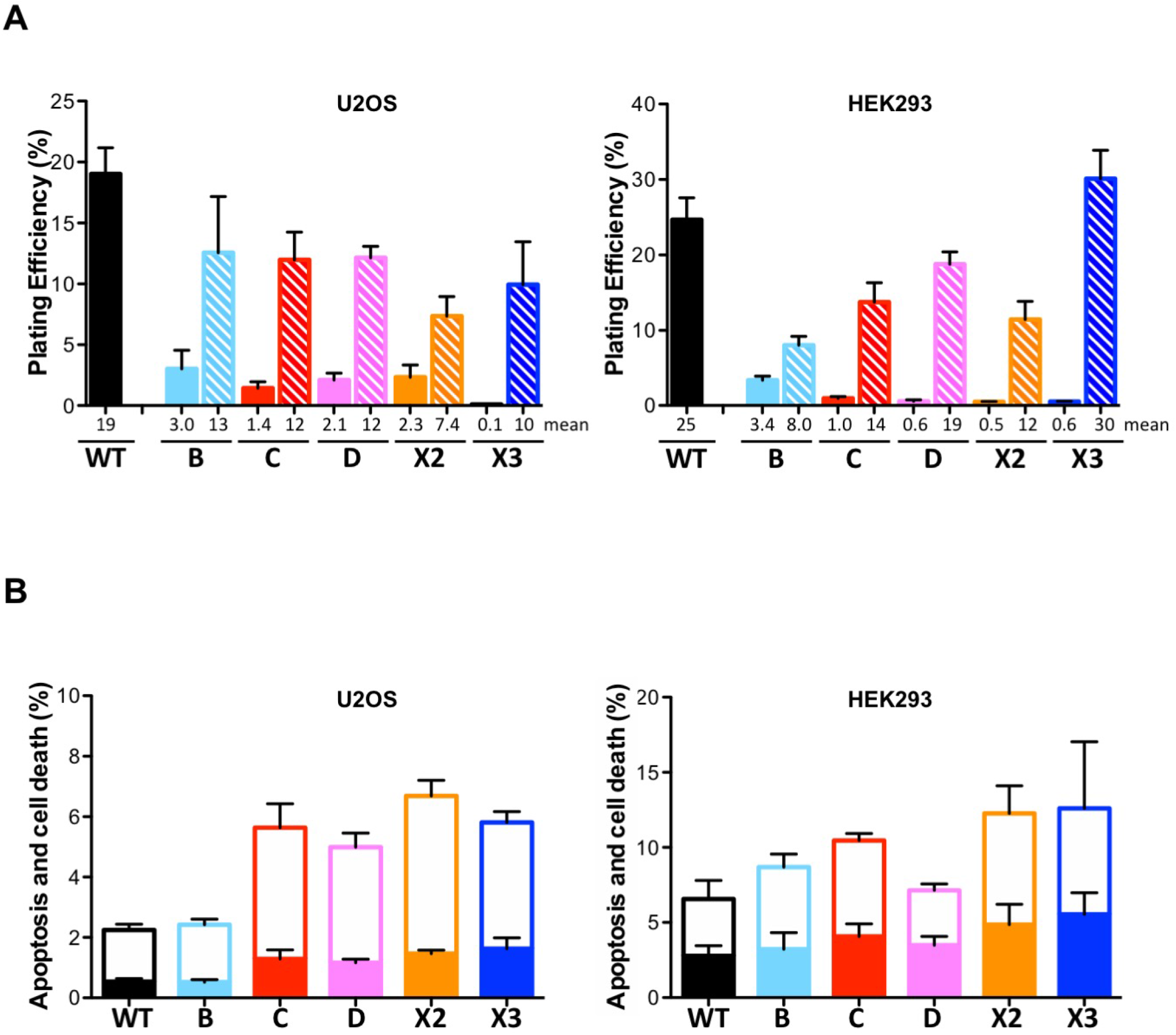
RAD51 paralog disruption in human U2OS and HEK293 cells leads to reduced plating efficiency but not to apoptosis. **(A)** Plating efficiency analysis was determined by colony formation assay normalized to initial seeding density. Filled and hatched bars are the mutant cells and the mutant cells stably complemented with a retroviral construct expressing the corresponding wild-type allele, respectively. Results are presented as means +/− SD from three experiments. **(B)** Apoptosis and cell death analysis using Annexin V and 7-Aminoactinomycin D staining of RAD51 paralog knockout cells. Open and filled bars represent fractions of dead cells and apoptotic cells, respectively. Results are presented as means +/− SD from three experiments.

### RAD51 paralog disrupted human cell lines are impaired in DNA double-strand break-induced homologous recombination

As the classical RAD51 paralogs have been implicated in the control of the RAD51 recombinase, the biological consequence of their disruption was examined by measuring HR levels. The conditional parents and derivative biallelic mutant RAD51 paralogs in MCF10A cells contain an integrated DR-GFP reporter to measure DSB-induced HR (Pierce et al. 1999; Feng and Jasin 2017) (Figure 3A). After Cre expression to generate mutants, we expressed I-SceI endonuclease using lentiviral transduction to induce a DSB in the DR-GFP reporter and monitored HR using FACS to quantify GFP+ cells. The RAD51 paralog disrupted cells have a ∼2 to >10-fold reduction in HR compared to the parental cells, with *RAD51B* disrupted cells showing the smallest reduction and *RAD51C* and *RAD51D* disrupted cells showing the largest reduction (Figure 3A).

**Figure 3.**
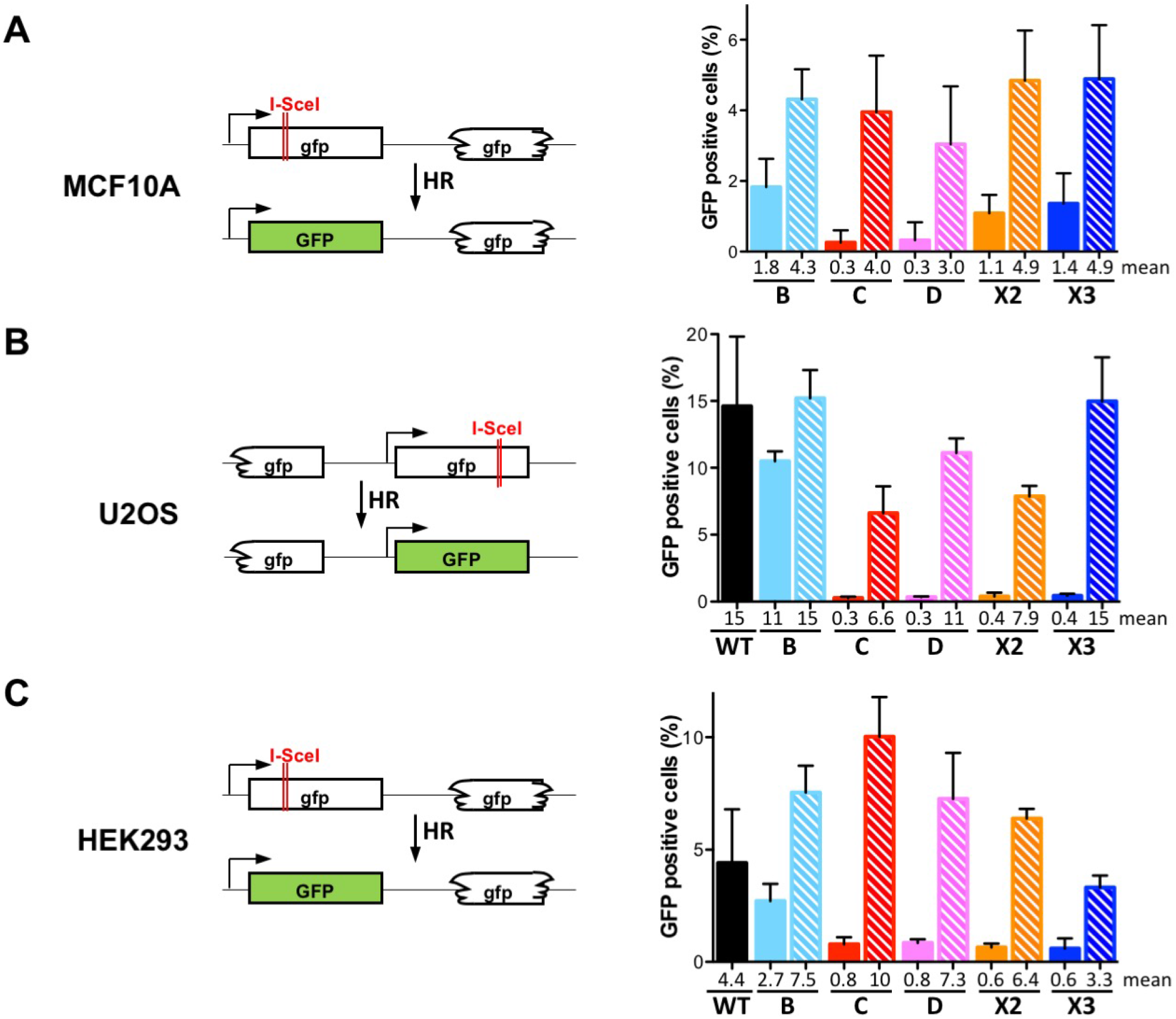
RAD51 paralog disruption lead to homologous recombination deficiency. **(A)** Schematic of the I-SceI-inducible direct repeat recombination reporter integrated in the genome of MCF10A (Pierce et al. 1999; Feng and Jasin 2017). Since RAD51 paralogs in MCF10A cells (except RAD51B) were inviable after Cre expression, the analysis was performed 2 days post-Cre when the mutant cells were still alive. Post-Cre mutant and parental cells were infected with I-SceI – expressing lenti-virus for 48 h. GFP+ cells were measured by FACS. Filled and hatched bars represent the mutant and parental cells, respectively. Data are presented as means +/− SD from at least three independent experiments. **(B, C)** Schematic of the I-SceI-inducible direct repeat recombination reporter integrated in the genome of U2OS (Puget et al. 2005) cells and HEK293 (Esashi et al. 2005). Cells were transfected with I-SceI-expressing plasmid and GFP+ cells were measured. Filled and hatched bars are for the mutant cells and the mutant cells stably complemented with a retroviral construct expressing the corresponding wild-type allele, respectively. Data are presented as means +/− SD from at least three independent experiments.

The U2OS and HEK293 cell lines also contain HR reporters integrated in the genome, DR-GFP in the case of HEK293 and a related HR substrate in U20S (Figure 3B and Figure 3C). Disruption of each RAD51 paralog significantly decreased HR efficiencies in comparison to wild-type cells in both cell backgrounds (Figure 3B and Figure 3C, filled bars). Of note, disruption of *RAD51B* resulted in only a moderate 1.5-fold reduction in HR efficiencies compared to wild-type cells, in contrast to disruption of the other RAD51 paralogs, which led to much greater reductions (up to nearly 40-fold). HR was rescued in mutant cells stably complemented with the corresponding wild-type RAD51 paralog alleles (Figure 3B and Figure 3C, hatched bars) confirming that all RAD51 paralogs are important for repair of DSBs by HR.

To determine whether there is any functional redundancy between the five classical RAD51 paralogs, transient overexpression of each was performed in the panel of mutant cell lines. As expected, overexpressing each paralog in the cell line lacking the cognate factor substantially restored HR efficiencies. In contrast, cross-complementation did not reverse the HR defect in any of the mutant cell strains (Figure S4). We conclude that each classical RAD51 paralog plays a specific function during HR.

In yeast, expression of Rad51 or Rad52 from a high-copy number plasmid partially suppresses the HR defect of strains deficient in the two Rad51 paralogs, Rad55 or Rad57 (Hays et al. 1995; Johnson and Symington 1995). Similarly, overexpression of RAD51 or RAD52 in classical RAD51 paralog deficient chicken cells partially suppresses their HR defects (Fujimori 2001; Takata et al. 2001). However, expression of either RAD52 or BRCA2 (under the control of the strong cytomegalovirus promoter) did not significantly reverse the defect in HR in any of the RAD51 paralog mutant cells (Figure S4). Of note, ectopic expression of RAD52 resulted in drastic reductions in HR, in both wild-type cells and *RAD51B* deficient cells (both U2OS and HEK293). Marginal enhancement of HR levels was observed by overexpression of RAD51 in U2OS cells but not in HEK293 cells (Figure S4). We conclude that the roles of the classical RAD51 paralogs are each distinct from that of BRCA2 or RAD52 and that increasing RAD51 levels is not sufficient to bypass HR deficiencies in cells deficient in any of the RAD51 paralogs.

### Stable RAD51 nuclear focus formation is impaired in classical RAD51 paralog disrupted U2OS cells

To explore the involvement of the classical RAD51 paralogs in RAD51 control, we next investigated spontaneous and IR-induced RAD51 nuclear foci formation in asynchronous populations by immunofluorescence in wild-type and RAD51 paralog deficient U2OS lines. Approximately 40% of wild-type cells were positive for spontaneous RAD51 focus formation (defined as cells with at least 5 foci/nucleus, Figure 4) in the absence of irradiation. In comparison, RAD51 paralog deficient cells exhibited reduced numbers of cells with RAD51 foci. RAD51 foci were apparent in ∼75 % of wild-type cells irradiated with 4 Gy, but were markedly reduced in all of the RAD51 mutant cells with *RAD51B* deficient cells exhibiting an intermediate phenotype. Defects in RAD51 foci formation were completely reversed by re-expressing each respective RAD51 paralog (Figure 4 hatched bars, and full data set in Figure S5A).

**Figure 4.**
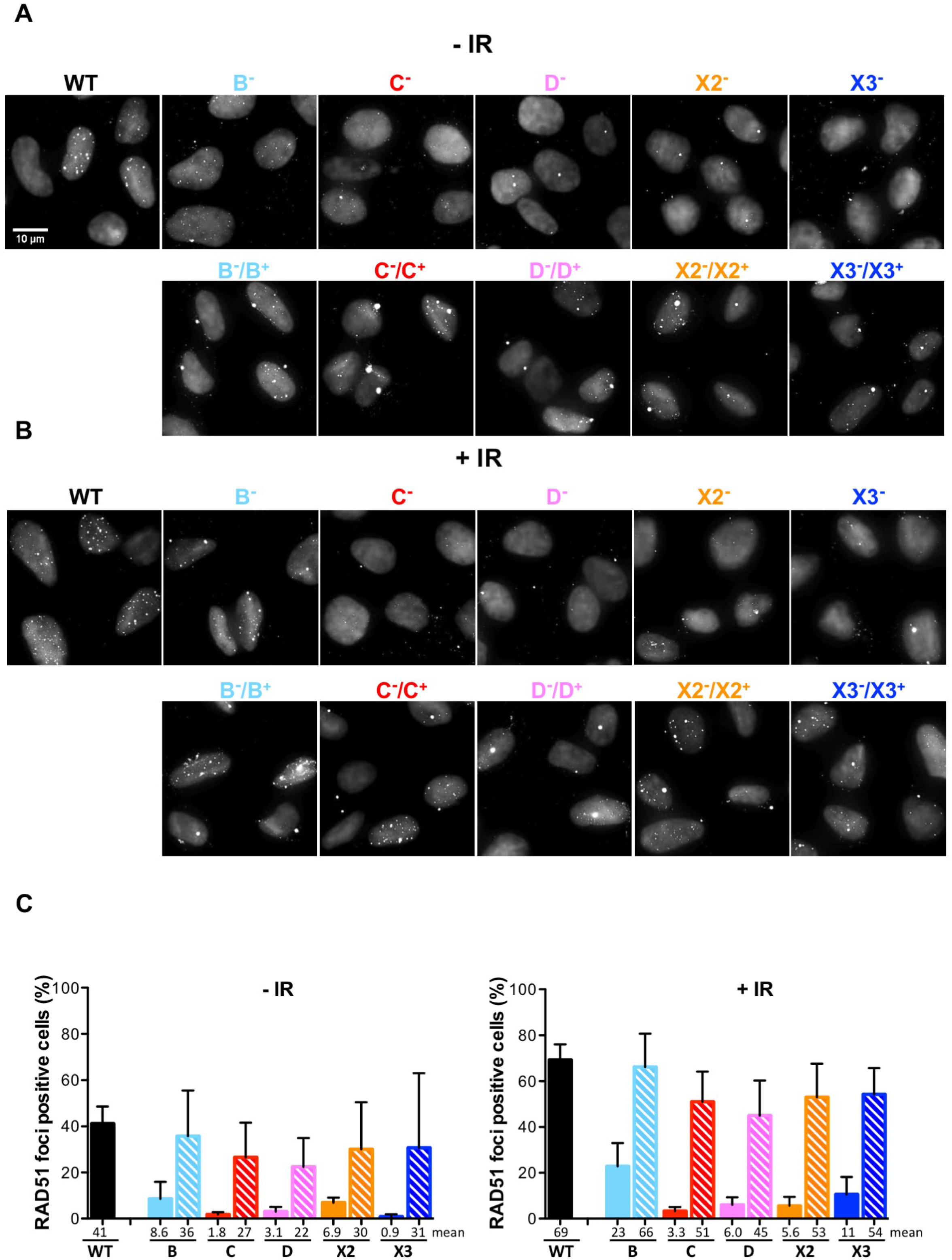
RAD51 nuclear focus formation is impaired in RAD51 paralog disrupted cells. **(A, B)** Detection by immunofluorescence of RAD51 nuclear foci in wild-type and RAD51 paralog disrupted U2OS cells, untreated (-IR) or 4 h after exposure to 4 Gy (+IR). Representative micrographs were obtained by confocal microscopy and shown as black and white overlays of RAD51 immuno-staining signal and DNA staining with DAPI to delineate nuclei. Top row panels are images obtained from wild-type and mutant cells. Bottom row panels are images obtained from the mutant lines stably complemented with a retroviral construct expressing the corresponding wild-type allele. Note that some RAD51 foci are detected outside of the nuclear region. These correspond to the accumulation of RAD51 at centrosomes (microtubule organizing centers) as verified by gamma-tubulin co-localization experiments (data not shown). **(C)** Quantification of RAD51 nuclear focus formation by counting cells with at least 5 foci per nucleus as positive. For wild-type and mutant cells, in two experiments at least 100 nuclei were scored, and at least 50 nuclei were scored in the third experiment. For the stably complemented mutant cells at least 50 nuclei were scored in three experiments. Data are presented as means +/− SD from three independent experiments. See Figure S5A for the entire data pools.

The function of XRCC3 in RAD51 foci formation is controversial (Chun et al. 2013; Yoshihara et al. 2004), and we investigated the impact of XRCC3 disruption on RAD51 foci formation more precisely over time. No increase in cells positive for RAD51 focus formation in *XRCC3* mutant cells compared to wild-type U2OS cells (Figure S5B) was observed at any time point. Similarly, no increase in cells positive for RAD51 focus formation was observed when RAD51C was overexpressed in *XRCC3* deficient cells (Figure S5C). We conclude that the classical RAD51 paralogs are crucial for formation of stable spontaneous and IR-induced RAD51 nuclear focus.

### Classical RAD51 paralog disruption sensitizes human U2OS and HEK293 cells to mitomycin C and olaparib

Mutations in RAD51C and XRCC2 result in Fanconi anemia, a syndrome associated with extreme sensitivity to drugs like mitomycin C (MMC) that induce DNA interstrand crosslinks (Park et al. 2016; Somyajit et al. 2010; Vaz et al. 2010b). MMC sensitivity of each U2OS mutant cell line was assessed by clonogenic survival assays. As these assays are challenging with cells that have reduced plating efficiency, conditioned medium was used which was found to enhance plating efficiency (Figure 5A). RAD51 paralog mutant cells were hypersensitive to MMC in comparison to wild-type cells (Figure 5B). However, *RAD51B* disrupted cells presented a less acute sensitivity to MMC than the other RAD51 paralog mutants. Moreover, the hypersensitivity to MMC was efficiently rescued in RAD51 paralog mutant cells stably complemented with the cognate wild-type RAD51 paralog genes (Figure S6A).

**Figure 5.**
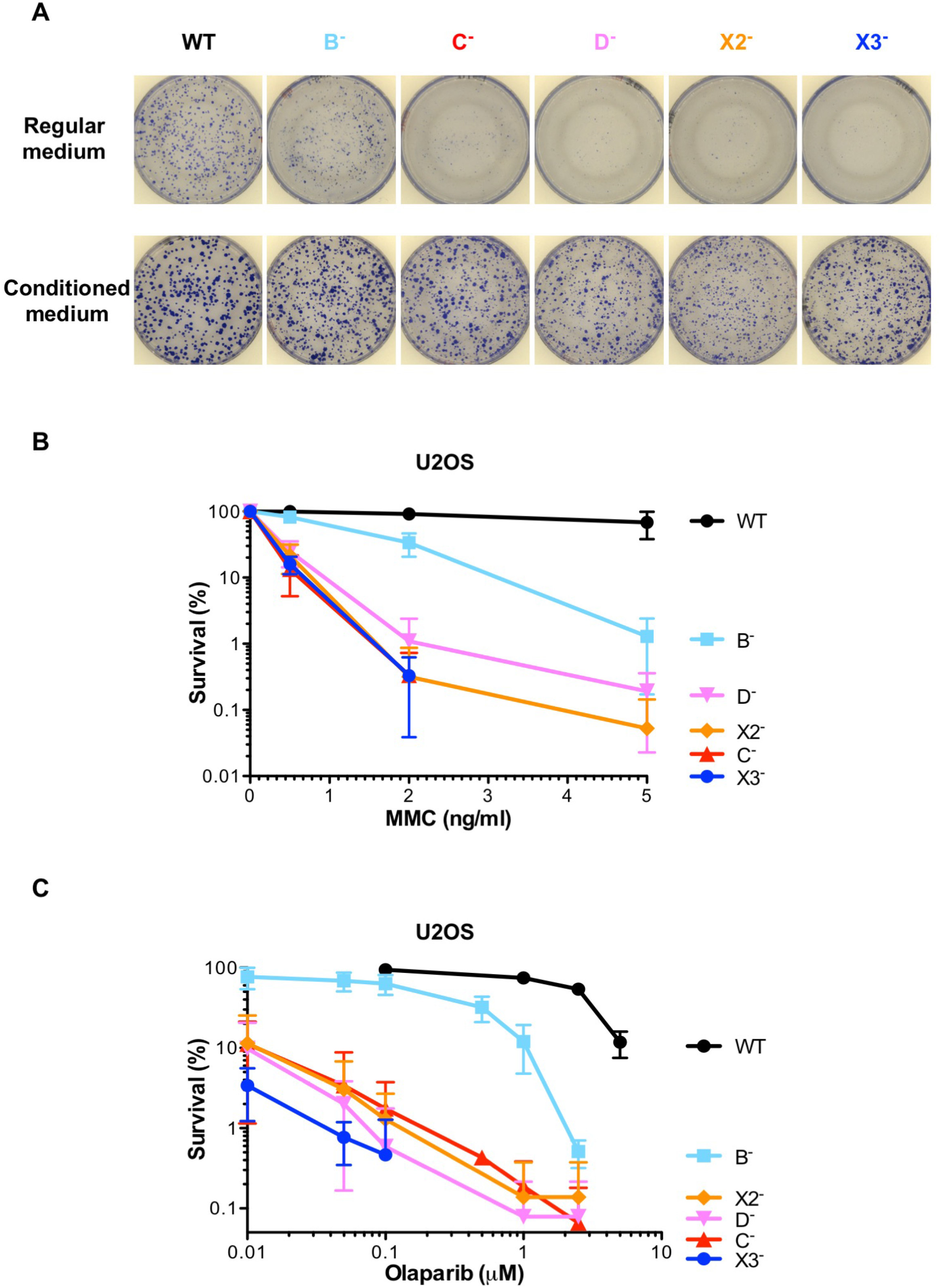
RAD51 paralog disruption sensitizes human U2OS cells to mitomycin C and olaparib. **(A)** Rescue of the plating efficiency defect of RAD51 paralog disrupted U2OS cells using conditioned medium. Images of the culture dishes stained with Coomassie to reveal colonies formed 7 to 10 days after seeding the wild-type and *RAD51B* mutant cells in regular versus conditioned medium, and after 14 to 15 days for the other mutants. **(B)** Survival curves obtained by clonogenic cell survival assays after treatment of exponentially growing U2OS cells with indicated doses of mitomycin C (MMC). Analyses of the mutant cells stably complemented with a retroviral construct expressing the corresponding wild-type allele are shown in Figure S6A. Results are presented as means +/− SD from at least three independent experiments. **(C)** Survival curves obtained by clonogenic cell survival assays after treatment of exponentially growing U2OS cells with indicated doses of olaparib. Analyses of the mutant cells stably complemented with a retroviral construct expressing the corresponding wild-type allele are shown in Figure S6B. Results are presented as means +/− SD from at least three independent experiments.

Poly(ADP-ribose) polymerase (PARP) inhibition is used in the clinic to kill HR-deficient tumor cells such as cells mutated for the breast cancer predisposition genes BRCA1 or BRCA2 by synthetic lethality (Bryant et al. 2005). Given the substantial HR deficiency of the RAD51 paralog knockout lines, we tested whether they could efficiently be killed by treatment with the clinically approved PARP inhibitor olaparib. RAD51 paralogs mutant cells were highly sensitive to olaparib in comparison to wild-type cells in clonogenic survival assays (Figure 5C). However, *RAD51B* knockout cells presented a less acute sensitivity to olaparib than the other RAD51 paralog mutants. Moreover, the hypersensitivity to olaparib was efficiently rescued in RAD51 paralog mutant cells stably complemented with the respective wild-type alleles (Figure S6B). Overall, the hypersensitivity to MMC and olaparib that we observed in U2OS cells with individual disruption of the classical RAD51 paralogs is consistent with their implication in protection and rescue of perturbed DNA replication forks.

## DISCUSSION

We report here the generation of viable individual knockouts of the five classical RAD51 paralogs in two different transformed human cell lines: U2OS cells of bone cancer origin and HEK293 cells that exhibit cancer stem cell features (Debeb et al. 2010). Furthermore, we also generated conditional mutants of each classical RAD51 paralog in non-transformed MCF10A mammary epithelial cells, and in this genetic background, only *RAD51B* disrupted cells are viable after Cre mediated excision of the complementing wild-type allele. It seems likely that the lethality associated with classical RAD51 paralog in MCF10A cells (with the exception of *RAD51B* disruption), but not in U2OS and HEK293 cells, is due to intact DNA damage sensing and response pathways in these non-transformed cells.

That RAD51B is distinct from the other RAD51 paralogs in terms of essentiality in MCF10A cells was surprising, given that it is a member of the BCDX2 complex, it can function with RAD51C to promote RAD51 function *in vitro*, and it is essential for embryogenesis in mice like the other members of this complex. In the last few years, genome-wide CRISPR-Cas9 genetic screens have been performed to determine gene essentiality in various human cell lines, each providing fitness scores to compare cells deficient in tested genes (Hart et al. 2015; Wang et al. 2015, 2017). We retrieved data from screens in 23 cell lines for each RAD51 paralog as well as for RAD52, which is not essential in mammalian cells (Rijkers et al. 1998; Sotiriou et al. 2016; Yasuhara et al. 2018). *RAD51B* gene-edited cells showed the best fitness scores in the vast majority of these cell lines compared to cells disrupted for any of the other four RAD51 paralogs (Figure S7), in line with our data that loss of *RAD51B* results in the weakest phenotype among the RAD51 paralogs. In fact, the *RAD51B* fitness scores are closer to those of *RAD52* than those of most of the other RAD51 paralogs. Interestingly, among the RAD51 paralogs, fitness scores for *XRCC2* gene-edited cells are intermediate between those of *RAD51B* and the three other RAD51 paralogs. By contrast, *RAD51D* fitness scores are generally among the worst of the RAD51 paralogs, including in KBM7 cells; however, in one study RAD51D was assessed to be non-essential in KBM7 cells (Blomen et al. 2015). Thus, although these large-scale screens can provide hints regarding fitness of mutants, a knockout strategy needs to be implemented to conclusively test for essentiality for a particular gene in a particular cell line.

Phenotypes, including defects in cell growth, HR, and sensitivity to MMC and olaparib, presented here for our collection of RAD51 paralog knockout lines, are overall similar to the ones observed in chicken DT40 cells (Takata et al. 2001). Consistent with the unique non-essentiality of RAD51B in MCF10A cells, *RAD51B* mutant human cell lines systematically showed the weakest phenotypes in all assays compared to the other RAD51 paralog mutants, a distinction not found in DT40 chicken RAD51 paralog knockout cells. A reason for this could relate to the apparent hyper-recombinogenicity of DT40 cell lines. Chinese hamster cells deficient for RAD51C, XRCC2 and XRCC3 have all been isolated in screens for IR sensitivity; RAD51B deficiency in hamster cells could also result in a relative milder phenotype and would therefore have escaped detection in IR sensitivity screens. Nevertheless, several studies suggested that the stability or steady-state levels of RAD51 paralog proteins are interdependent (Lio et al. 2004; Chun et al. 2013; Gildemeister et al. 2009; Katsura et al. 2009; Rodrigue et al. 2006). For instance, RAD51C deficiency was reported to result in decrease cellular levels of RAD51B, RAD51D and XRCC3. A possibility could be that *RAD51B* disruption does not impact on the levels of the other RAD51 paralogs and thus causes a less profound phenotypic impact. Alternatively, RAD51B could play a regulatory role rather than being directly involved in controlling RAD51 dynamics.

While the U2OS and HEK293 RAD51 paralog mutant lines reported here are viable, they are less fit than their wild-type counterparts with a tendency to accumulate dead cells in culture. Determining the reasons for the slow growth of the mutant lines will require further analysis. They will likely be linked to the role of the RAD51 paralogs in DNA replication fork maintenance given their implication in repair of collapsed replication forks by HR and protection of stalled replication forks (Somyajit et al. 2015; Saxena et al. 2018). Defects in replication fork maintenance could lead to cell cycle progression anomalies. In this context, the studies suggesting direct roles of some of the classical RAD51 paralogs in cell cycle checkpoint signaling are intriguing (Badie et al. 2009). However, the exact roles of the classical RAD51 paralogs in these signaling mechanisms are still unresolved, but the reagents generated in this study will help address some of these issues. Altogether, the impact on cell growth and physiology of individual disruption of the five classic RAD51 paralogs in human U2OS and HEK293 cells parallel the ones reported for the individual RAD51 paralog knockouts in chicken DT40 cells (Takata et al. 2000, 2001). Given that DT40 cells are derived from a bursal lymphoma induced by avian leukosis virus infection, and are therefore of cancer origin, our results suggest that divergent DNA repair processes operate in cancer cells compared to normal cells.

Defects in spontaneaous and IR-induced RAD51 nuclear focus formation were observed in all of the RAD51 paralog mutant U2OS lines, indicating that all five classical RAD51 paralogs are required for stable RAD51 nuclear focus formation. These results contrast with a previous report in which U2OS and HCT116 *XRCC3*-deficient cells showed normal formation of RAD51 nuclear foci in response to IR (Chun et al. 2013). However, the latter study in U2OS cells was based on siRNA-mediated depletion of XRCC3 rather than loss-of-function alleles. The results from HCT116 cells remain puzzling, although it should be noted that HCT116 cells, which are of colon cancer origin, are mismatch repair deficient and have accumulated numerous additional mutations (Bhattacharyya et al. 1994). Nevertheless, the results from our study suggest that all classical RAD51 paralogs are required for detection of stable RAD51 containing repair complexes, with RAD51B having a more modest role. We propose that a collaborative dynamic interplay between all five classical RAD51 paralogs is involved in controlling stabilization and remodeling of RAD51 complexes at DNA repair sites, for example, as proposed for the yeast RAD51 paralog complex Rad55-Rad57 and C. elegans RAD51 paralog complex RFS-1/RIP-1 (Liu et al. 2011; Taylor et al. 2015).

In conclusion, we have generated three sets of human isogenic lines with individual disruption of the five classical RAD51 paralogs that can now be used by the community to further explore their cellular functions. The mutant lines harbor a HR reporter stably integrated in their genome, which provides a means to rapidly analyze the impact of patient-derived mutations by functional complementation. As a proof of concept, in this study we have identified five highly conserved residues in RAD51B to be mutated in tumors from MSK-IMPACT database (Figure S8A). Three of these are located in the Walker domains. Yeast 2-hybrid analysis on RAD51B G108D, V207R, and G341V showed reduced interactions with RAD51C while RAD51B K114Q and R159H showed wild-type like interactions with RAD51C (Figure S8B). We overexpressed these mutant RAD51B proteins in *RAD51B^-/-^* in HEK293 cells and performed HR experiments. RAD51B K114Q and R159H were able to complement the HR defect of *RAD51B^-/-^* cells like the wild-type RAD51B but RAD51B G108D, V207R, and G341V behaved like null mutants (Figure S8C). These results suggest that Walker A and B domains of RAD51B are important for its interaction with RAD51C and for its function in HR. Mutating the highly conserved lysine (K114) in the Walker A domain to glutamine (Q) in RAD51B did not affect its complementing activity in HR assays suggesting that either this lysine residue is not as important as the other conserved residues in this domain or that the substitution of the lysine to glutamine would be tolerated by cells. In addition, we have recently utilized these cell lines to perform similar HR analysis with RAD51C and RAD51D patient-derived reversions and mutations, respectively (Baldock et al. 2019; Kondrashova et al. 2017). This kind of analysis will be particularly useful for investigation of the numerous RAD51 paralog variants of unknown significance found in patient cancer cells which may in the long-term help optimize personalized cancer therapies. Moreover, the cell lines will also be valuable for clinically oriented studies aimed at understanding drug resistance and synthetic lethality mechanisms. These reagents will be helpful for in depth fundamental analysis of the roles of the classical RAD51 paralogs in genome maintenance mechanisms, including, not only HR-related mechanisms, but also DNA damage cell cycle checkpoint, chromosome segregation and other functions.

## MATERIALS AND METHODS

### Cell culture

The MCF10A cell line containing an integrated DR-GFP reporter (Feng and Jasin 2017) was grown in DMEM HG/F-12 supplemented with 5% horse serum, 1% penicillin and streptomycin, 100 ng/mL cholera toxin, 20 ng/mL epidermal growth factor, 0.01 mg/mL insulin, and 500 ng/mL hydrocortisone in a 5% CO_2_ atmosphere at 37°C. Infection of MCF10A cells with Lenti-Cre was performed as described in (Feng and Jasin 2017). The HEK293 DR-GFP cell line (Esashi et al. 2005) and the U2OS SCR#18 cell line kindly provided by Dr. Ralph Scully and Dr. Nadine Puget (Puget et al. 2005) were grown in DMEM supplemented with 10% fetal bovine serum, 1% penicillin and streptomycin in a 5% CO_2_ atmosphere at 37°C. For HEK293 DR-GFP cells, plates were pre-coated with poly-L-lysine (Sigma). All the cell lines were tested mycoplasma negative.

### Establishment of conditional RAD51 paralogs and knockouts in MCF10A Cells

To generate the five conditional RAD51 paralog cell lines, MCF10A cells were co-transfected with a donor plasmid (see Table S6) that contained a full-length cDNA for each paralog driven by the CAG promoter (cytomegalovirus enhancer fused to the chicken beta-actin promoter) and flanked by LoxP sites and vectors expressing TALENs toward AAVS1 locus. Transfections were performed using electroporation (Gene Pulser II, Bio-Rad; 350V, 1000µF) and post-transfected cells were selected with G418 (0.2 mg/ml) for a week. G418 resistant colonies were picked and analyzed for the presence of cDNA using cDNA specific primers. Once all five conditional RAD51 paralogs lines were generated, either CRISPR-Cas9 or TALENs were used to disrupt endogenous alleles as described below.

To generate *RAD51B* knockouts paired gRNA were used. The oligonucleotides used to generate the guides are shown in Figure S1. To knockout *RAD51C*, *RAD51D*, *XRCC2* and *XRCC3*, TALENs were used. Left and right TALEN recognition sequences are shown in Figure S1. After transfection of either gRNA or TALENs, individual colonies were picked, expanded and PCR was performed across the exon of interest with the primer pairs described in the Table S2. For *RAD51B*, paired gRNA technique was employed, and the PCR products were fractionated on a 2.4% gel to determine size difference. The clones that contained shorter PCR products were further analyzed by TOPO cloning the PCR products and DNA sequencing. At least ten colonies were sequenced to determine the indel type. For the other four RAD51 paralogs, PCR was performed using the primer pair described in Table S2 across the exon of interest. This PCR product was digested with the indicated restriction enzyme (Table S2). If the PCR product was completely digested, the clone would be genotyped as wild-type, however if the PCR product was resistant to the indicated restriction enzyme, the clone would be identified as a mutant. These mutant clones were further analyzed by TOPO cloning the PCR products and DNA sequencing. At least ten colonies were sequenced to determine the indel type.

### Establishment of RAD51 paralogs U2OS and HEK293 knockout cell lines

The RAD51 paralogs knockout cell lines were generated using the *S. pyogenes* CRISPR-Cas9 genome editing system (Ran et al. 2013). Oligonucleotides used to generate vectors expressing guide RNAs targeting *RAD51B*, *RAD51C*, *RAD51D*, *XRCC2* and *XRCC3* genes, and oligonucleotides for PCR amplification of the targeted genomic locus are listed in Table S3 and Table S4, respectively. Guide RNA sequences were annealed and cloned into BbsI sites of pSpCas9(BB)-2A-GFP vector (Addgene plasmid #48138, pX458). These pX458-derived plasmids were transfected into U2OS and HEK293 cells using polyethyleneimine (PEI) at 2 µg PEI/µg DNA. GFP+ cells were sorted in 96-well plates 48 h post transfection using a FACS Aria III cell sorter (BD BioSciences). Single cells were grown until formation of viable individual clone in DMEM medium supplemented with 50% fetal bovine serum and 1% penicillin/streptomycin. Cells were expanded and knockout clones were confirmed by western blotting and sequencing. Genomic DNA was isolated from edited clones and non-edited control cells using the DNeasy Blood & Tissue Kit (Qiagen). RAD51 paralog specific loci were PCR amplified using gene specific PCR primers Table S4. The PCR products were cloned into the pCRBlunt vector and transformed in Top10 *E. coli* bacteria. Sanger sequencing was used to analyzed at least seventeen individually cloned amplicons.

### Cell extracts and Immunoblotting

Cell lysates were prepared by resuspending cell pellets in RIPA buffer (150 mM NaCl, 50 mM Tris pH 8.0, 5 mM EDTA, 0.5% sodium deoxycholate, 0.1% SDS, 1.0% Nonidet P-40, 2.0 mM phenylmethylsulfonyl fluoride, 1 mM Na_3_VO_4_) supplemented with protease and phosphatase inhibitors (Pierce, Life Technologies). The lysates were incubated on ice for 30 min and cleared by centrifugation (14,000 rpm for 30 min at 4°C). Protein concentrations were measured using Bradford Biorad Protein Assay kit. Equal amount of protein extract (50-100 µg) was resolved on Bolt™ 4-12% Bis-Tris Plus Gels (Invitrogen) and transferred onto PVDF membrane. Membranes were blocked for 1 h using 3% BSA in TBST (50 mM Tris-HCl pH 8, 150 mM NaCl, 0.1% tween-20). Proteins were detected using the primary antibodies listed in Table S6. Secondary antibody detection was performed using HRP-conjugated goat anti-rabbit (1:5000), or goat anti-mouse (1:5000) (Dako). Immunoblots were developed using Chemiluminescent ECL western blotting detection reagents (ECL™Prime Western Blotting System, RPN2232 or ECL Select™ Western Blotting Detection Reagent RPN2235 from GE Healthcare). Images were analyzed on a ChemiDocMP system (Biorad). Loading control and normalization were assessed using anti-GRB2 antibodies.

### Doubling time

Doubling times were measured using the CellTiter-Glo Luminescent Cell Viability Assay (Promega) using manufacturer recommendation. Briefly, 5000 cells were seeded in 96-well plates and measurements were performed initially and 24, 48, 72 and 96 h. Cell doubling time was calculated using GrahPad Prism software by nonlinear regression (exponential growth equation) analysis.

### Plating efficiency

Plating efficiency was determined by colony formation assay. Cells were seeded in 6-well plates at 500 and 2500 cells per well and grown for 7 to 15 days. Cells were washed with PBS, fixed and stained with 50% ethanol, 7% Acetic acid, 1 g/L Coomassie blue R250. Colonies containing more than 50 cells were counted. Plating efficiencies were calculated as number of colonies/number of plated cells x 100.

### Apoptosis assays

Apoptosis was measured using Annexin V and 7-Aminoactinomycin D labeling (eBioscience). Cells were seeded onto 6-well plates. After 24 h, exponentially growing cells were collected and washed with cold PBS. Then, resuspended cells were stained at 10^6^ cells/ml in 100 µl of 1X Annexin V Binding Buffer containing Annexin V and 7-Aminoactinomycin D according to the manufacturer’s instructions (eBioscience). After incubation for 15 minutes at room temperature in the dark, 100 µl of 1X Annexin V Binding Buffer were added and cells were analyzed by FACS. The percentage of viable cells (low Annexin V/low 7-Aminoactinomycin D) was determined.

### Genotyping of MCF10A cells post-Cre treatment

Post-Cre MCF10A cells were seeded at a density of 1000-2000 cells / 10 cm plate in duplicates and allowed to grow for 11 days. After 11 days, one set of plates were washed with PBS, fixed with methanol for 30 min and stained with Giemsa (620G-75, EMD Millipore). Colonies were picked from second set of 10 cm plates. Genomic DNA was extracted from these colonies and PCR was performed across *AAVS1* locus using the following primers to confirm excision of RAD51 paralog. Forward primer: 5’-ATTGTGCTGTCTCATCATTTTGGC and Reverse primer: 5’-CTGGGATACCCCGAAGAGTG. PCR product sizes before and after Cre treatment are: ∼2.5 kb and ∼1.3 kb respectively.

### Clonogenic survival assays

U2OS cells were seeded in conditioned medium in triplicate in 6 cm plates and allowed to attach for 4 h before treatment with mitomycin C (Roche 10 107 409 001) or olaparib (Selleckchem AZD2281, Ku-0059436) at the indicated doses. Conditioned medium was obtained by mixing 50% fresh DMEM supplemented with 10% FBS and 1% penicillin/streptomycin and 50% filtered medium obtained from U2OS cells grown for 48 h and supplemented with 1% L-glutamate (Life Technologies). After 10 to 14 days, cells were washed, fixed and stained with 50% ethanol, 7% Acetic acid, 1 g/L Coomassie blue R250. Colonies containing more than 50 cells were counted. Results were normalized to untreated cells. For each genotype, cell viability of untreated cells was defined as 100%.

### DR-GFP Homologous Recombination assays

Post-Cre MCF10A cells were infected with I-Sce-I expressing lentivirus, cells were washed 24 h later and HR was measured 48 h later by quantifying the percentage of GFP+ cells by FACS (Becton Dickinson FACScan). U2OS and HEK293 cells were seeded into 10 cm plates 24 h prior to transfection with 2.5 µg of the I-SceI expression vector pCBA-SceI (Addgene plasmid # 26477, Richardson et al. 1998) or an empty vector in combination with 0.5 µg of pcDNA-RFP using polyethylenimine (PEI) at a ratio of 2:1 of PEI:total DNA. Cells were collected 72 h post-transfection and multicolor FACS analysis was performed with a LSRII apparatus (Becton Dickinson). Data were analyzed with FlowJo software (Tree Star, Ashland). RFP signal was used as transfection efficiency control and double positive cells (GFP+/RFP+) were counted as positive for HR events. Results were represented as a ratio of double-positive cells to the total number of RFP-positive cells. For transient complementation assay, 2 µg of pCMV plasmids expressing RAD51 paralogs, RAD51, RAD52 or BRCA2 were transfected in combination with 2.5 µg of pCBASceI and 0.5 µg of pcDNA-RFP. For stable complementation assays, cells transfected with retroviruses expressing RAD51 paralogs were used. For RAD51B cancer-associated mutant analysis, *RAD51B^-/-^* HEK293 cells were transfected with 1 µg of the I-SceI expression vector pCBA-SceI and 1 µg of the RAD51B expression vector. Cells were collected 48 h post-transfection and FACS analysis was performed by flow cytometry (BD FACScan), and data were analyzed using FlowJo software.

### Plasmid constructs

Human RAD51 paralog cDNAs were cloned into NcoI/XbaI sites of pCMV-myc-nuc (ThermoFisher). Human RAD51 cDNA was cloned into the NcoI/SalI sites of pCMV-myc-nuc. Human RAD52 cDNA was cloned into the NcoI/XhoI sites of pCMV-myc-nuc. Human BRCA2 cDNA was cloned into the NcoI/XhoI sites of a modified pCMV-myc-nuc. For the retroviral constructs expressing the wild-type RAD51 paralogs, FLAG tagged-human RAD51 paralog cDNAs were PCR amplified to add EcoRI-kozak-start-flag sequences to the 5’ end of each RAD51 paralog as indicated in Table S6. Each RAD51 paralog was then cloned into EcoRI/SalI sites of the pWZL-hygro retroviral vector (Addgene plasmid #18750). Retroviruses were produced in Phoenix-AMPHO cells (ATCC® CRL-3213™) and harvested 48 h after transfection. RAD51 paralog disrupted cells were transduced with retroviruses in presence of 2 µg/mL of polybrene (Sigma) and selected 72 h post-transduction with 300 μg/ml (U2OS) or 200 μg/ml (HEK293) hygromycin B for 7 days. RAD51B point mutants were made in yeast 2-hybrid plasmids (pGAD-C1 and pGBD-C1) and a mammalian expression plasmid (pCMV) using site-directed mutagenesis. The oligonucleotides used to generate these mutants are listed in Table S6. Yeast 2-hybrid experiments were performed as previously described (Kondrashova et al. 2017). All plasmids are listed and described in Table S6.

### Immunofluorescence

For RAD51 nuclear focus formation analysis, cells were cultured on coverslips for 24 h and then irradiated or not with 4 Gy using a RS2000 generator (Radsource). After irradiation, cells were allowed to recover for 4 h. Cells were then washed with PBS, treated with CSK buffer (10 mM PIPES pH 7.0, 100 mM NaCl, 300 mM sucrose, 3 mM MgCl_2_, 0.7% Triton X-100) for 2 min at 4°C, washed with PBS and fixed with 4% paraformaldehyde for 10 min at 4°C before an additional fixation step of 2 min with glacial methanol. Coverslips were rinsed with PBS and blocked for 1 h in PBS containing 0.1% Triton X-100 and 5% BSA. Cells were stained using a rabbit anti-RAD51 serum (1:5000) (Essers et al. 2002) in 0.5% BSA PBS at room temperature for 1 h prior to incubation with AlexaFluor 594 or 488 anti-rabbit secondary antibodies (1:5000, Molecular Probes) in 0.5% BSA PBS. Coverslips were mounted onto slides with Vectashield mounting media containing DAPI (Vector laboratories) and images were obtained using a Zeiss Axio Imager Z2 microscope with a 63x oil immersion objective. Maximum-intensity projection images were generated to display foci in all sections and were analyzed using ImageJ software. Automatic counting was performed using FoCo (Lapytsko et al. 2015) and validated manually.

### Statistical Analysis

Statistical analysis was performed using GraphPadPrism version 6.0 (GraphPad Software). Measurements are presented as means ± SD. Comparisons between two groups were analyzed by performing unpaired one-way ANOVA followed by Tukey’s test. A base p value of < 0.05 was considered statistically significant.

### Analyses of mycoplasma and viral contaminations

To verify that cell lines were free of mycoplasma or viral contaminations, mycoplasma contamination was monitored as described (Uphoff and Drexler 2014). Detection of EBV, HBV, HCV, HIV-1, HIV-2, HTLV-I/II, MLV, and SMRV was carried out as described previously (Uphoff et al. 2010, 2015).

### Authentication of cell lines and confirmation of targeted gene disruptions

For authentication of cell lines, genomic DNA was isolated using the High Pure PCR Template Preparation Kit (Roche Life Science). STR DNA genotyping at DSMZ was carried out as described previously using a nonaplex PCR reaction of eight highly polymorphic STR loci plus Amelogenin-based gender determination (Dirks and Drexler 2013). Generated STR profiles have been compared with the international STR Reference Database of DSMZ (Dirks et al. 2010). All derivatives of MCF-10A, U2OS and HEK293 showed full authenticity.

DSMZ performed an independent confirmation of gene disruptions in U2OS SCR#18 and HEK293-DR-GFP derivatives specific for RAD51B, RAD51C, RAD51D, XRCC2, and XRCC3 disruptions as well as their reconstituted counterparts. Using primers presented in Table S4, targeted DNA regions were amplified and respective amplicons cloned into pGEM-T (Promega) and the products subjected to Sanger sequencing. All disruptions of RAD51B, RAD51C, RAD51D, XRCC2 and XRCC3 could be confirmed.

### Quantitative PCR

To verify overexpression of RAD51B, RAD51C, RAD51D, XRCC2, and XRCC3 in complemented cell lines, total RNA was isolated using TRIzol reagent (Invitrogen) according to the manufacturer’s instructions. cDNA was synthesized from 5 µg of RNA with random primers using Superscript II Kit (Invitrogen, Thermo Fisher). Real-time quantitative PCR was performed with Taqman probes from Applied Biosystems (Hs01568763_m1 for RAD51B, Hs04194939_s1 for RAD51C, Hs00979545_g1 for RAD51D, Hs03044154_m1 for XRCC2, Hs00193725_m1 for XRCC3, 4333769F for TBP) and the Taqman Fast Advanced Master-Mix (Applied Biosystems, Thermo Fisher) using the 7500 Fast Real-Time PCR System (Applied Biosystems). Relative expression was evaluated using the ΔΔCt-method and TBP as endogenous control.

## ACKNOWLEDGMENTS

We thank Rémy Castellano and Audrey Restouin from the CRCM Target platform for help with cellular tests; Davide Normanno for help with microscopy; Marine Charpentier for TALEN plasmid construction; Jun Huang, Paul Russell, David Schild, Roland Kanaar and Kevin Hiom for sharing reagents; Sneha Saxena and Ganesh Nagaraju for advice with XRCC2 detection by immunoblotting; and Katheryn Meek and Bertrand Llorente for critical reading of the manuscript.

This work was supported by CRCM state subsidies from CNRS, Inserm, Institute Paoli-Calmettes and Aix-Marseille University; grant ANR-II-INSB-0014 (J-P.C., C.G.); MSK Cancer Center Support Core Grant (NIH P30CA008748); NIH Grants F32GM110978 (R.P.), R35GM118175 (M.J.), R01CA185660 (M.J.), F31ES027321 (M.R.S.), ES024872 (K.A.B.); American Cancer Society Grant 129182-RSG-16-043-01-DMC (K.A.B.); Stand Up to Cancer Grants SU2C-AACR-IRG-02-16 (K.A.B.) and SU2C-AACR-DT16-15 (M.J.); Postdoctoral fellowships from the French National League against Cancer and from Aix-Marseille University Foundation (E.B.G.); and the French National League against Cancer (M.M.)

## AUTHOR CONTRIBUTIONS

Conceptualization, E.B.G., S.G., R.P., M.J., M.M.; Investigation, E.B.G., S.G., R.P., G.J.B.; Writing – Original Draft, E.B.G., S.G., R.P.; Writing – Review & Editing, E.B.G., S.G., R.P., M.J., M.M.; Resources, M.R.S., K.A.B., A.D.C., J.P.C., C.G.; Validation, W.G.D., S.E.; Funding Acquisition, R.P., M.J., M.M.; Supervision, R.P., M.J., M.M.

## AVAILABILITY OF REAGENTS

U2OS and HEK293 derivatives including wild-type, mutant and complemented mutant lines have been deposited at the Leibniz Institute DSMZ, German Collection of Microorganisms and Cell Culture and the MCF10A mutant lines will be sent to this resource as well for future distribution.

---

**Table S1.** Designation of mutant clones.

|  | Gene |  |  |  |  |
| --- | --- | --- | --- | --- | --- |
|  | *RAD51B*<br>Chromosome 14<br>Forward strand<br>Transcript RAD51B-205<br>ENST00000471583.5 | *RAD51C*<br>Chromosome 17<br>Forward strand<br>Transcript RAD51C-201<br>ENST00000337432.8 | *RAD51D*<br>Chromosome 17<br>Reverse strand<br>Transcript RAD51D-202<br>ENST00000345365.10 | *XRCC2*<br>Chromosome 7<br>Reverse strand<br>Transcript XRCC2-201<br>ENST00000359321.1 | *XRCC3*<br>Chromosome 14<br>Reverse strand<br>Transcript XRCC3-201<br>ENST00000352127.11 |
| Cell line |  |  |  |  |  |
| MCF10A (parental) | B1 | C15 | D8.3 | X2-12 | X3-21 |
| MCF10A (mutant) | B-Δ87 | C-Δ16Δ41 | D-Δ10Δ19 | X2-Δ7Δ4 | X3-Δ42 |
| U2OS | B-8 | C-15 | D-4 | X2-5E | X3-6A |
| HEK293 | B-9 | C-2 | D-16 | X2-13 | X3-5 |

**Table S2.**
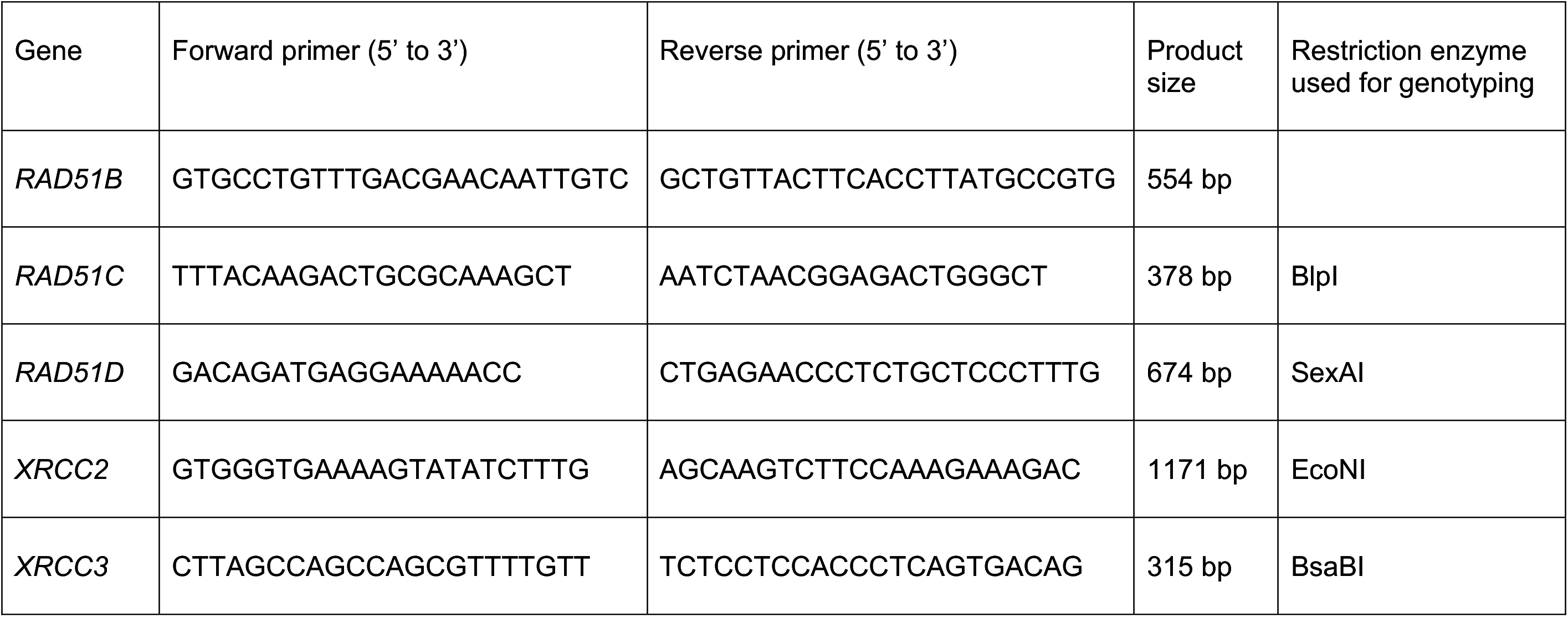
Genomic PCR primers for MCF10A cells.

**Table S3.**
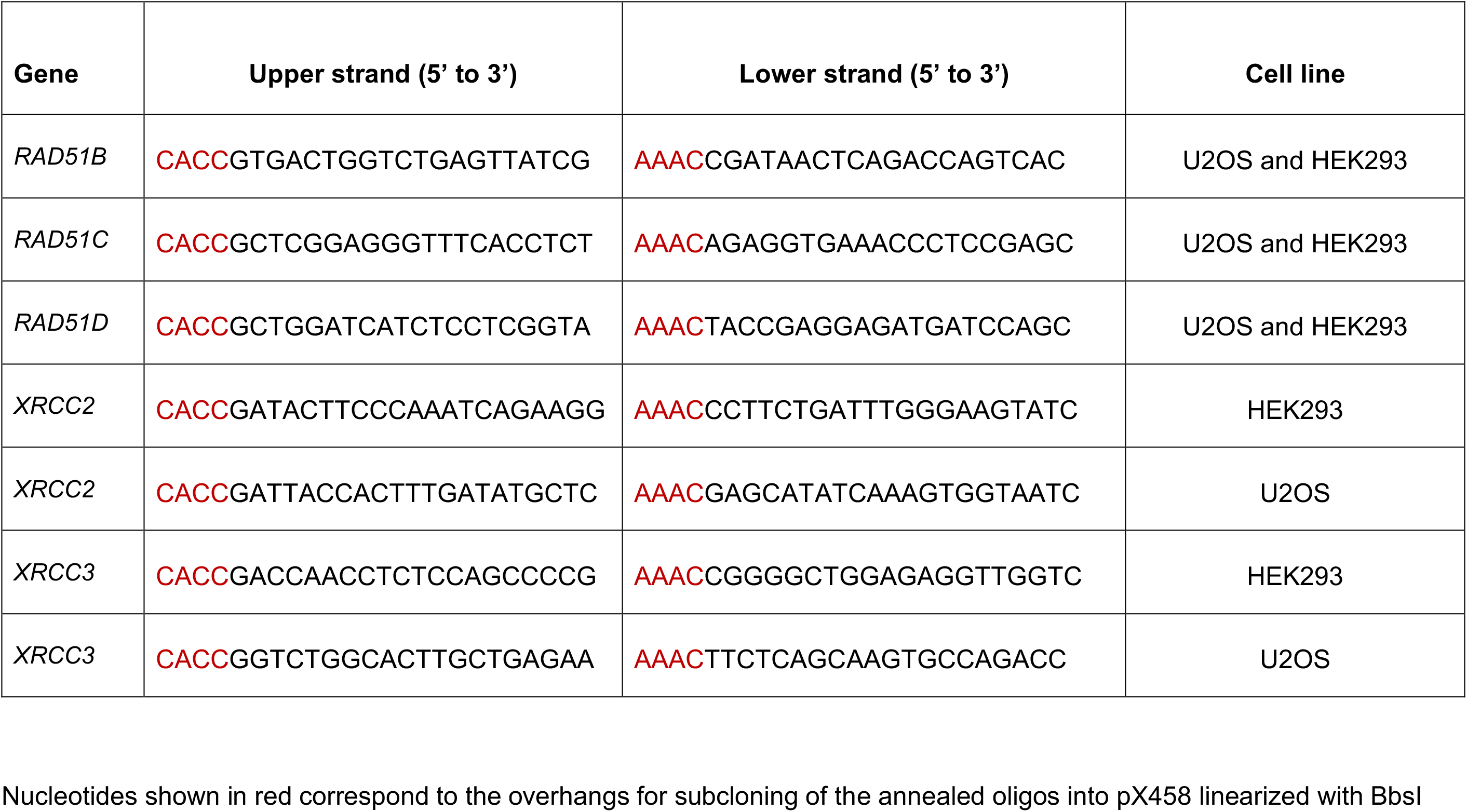
Oligonucleotides for gRNAs targeting RAD51 paralogs.

**Table S4.**
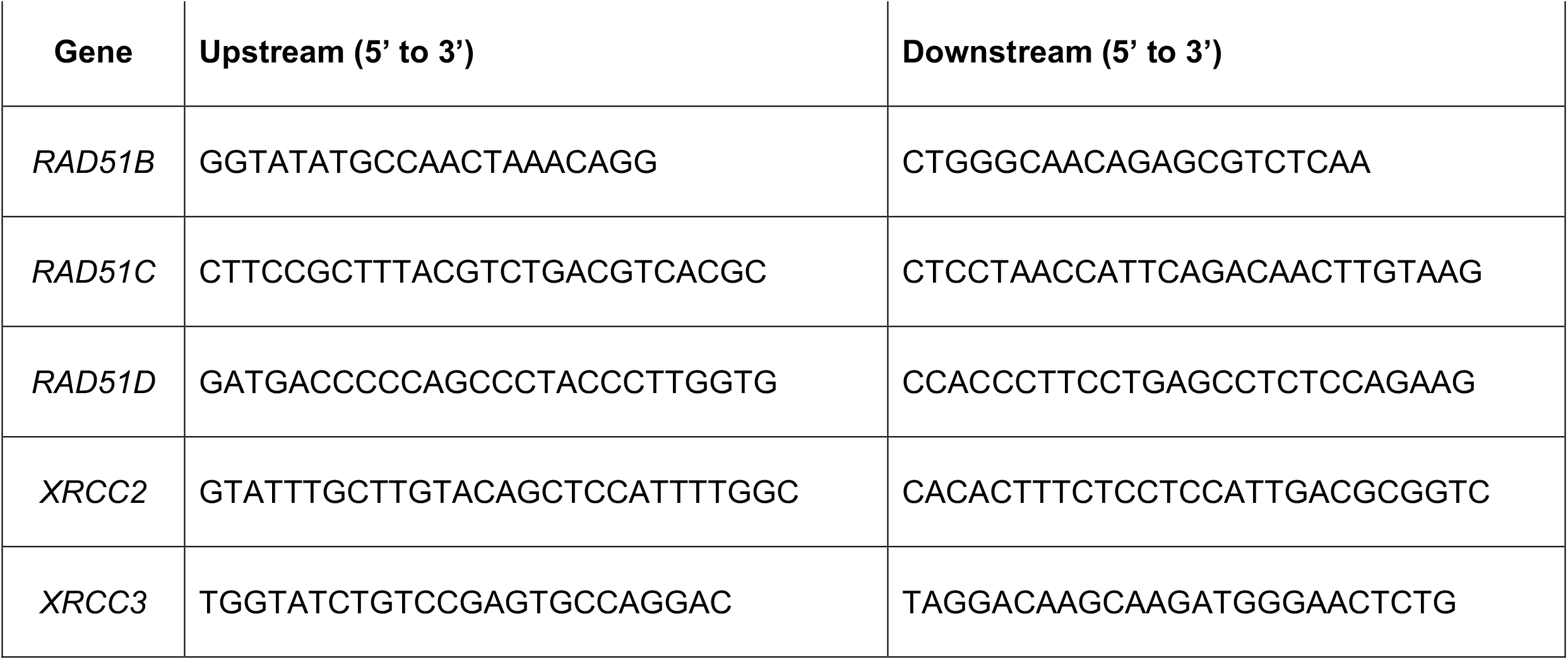
Genomic PCR primers for U2OS and HEK293 cells.

**Table S5.** Antibodies used in this study.

| Protein | Source | Reference | Dilution for WB or IF | Antigen residues |
| --- | --- | --- | --- | --- |
| RAD51B | SantaCruz | sc-377192 | 1/200 (WB) | Residues 141-255 |
| RAD51C | Abcam | ab95069 | 1/2000 (WB) | Residues 326-376 of hRAD51C (NP_478123.1) |
| RAD51D | Abcam | ab202063 | 1/1000 (WB) | Residues 1-100 |
| XRCC2 | SantaCruz | sc-365854 | 1/200 (WB) | Residues 6-33 |
| XRCC3 | SantaCruz | sc-271714 | 1/200 (WB) | Residues 1-300 |
| GRB2 | BD Bioscience | 610112 | 1/5000 (WB) | Residues 1-217 |
| RAD51 | Roland Kanaar | batch 2307<br>(Essers et al. 2002) | 1/5000 (IF) | Full-length RAD51 |

**Table S6.**
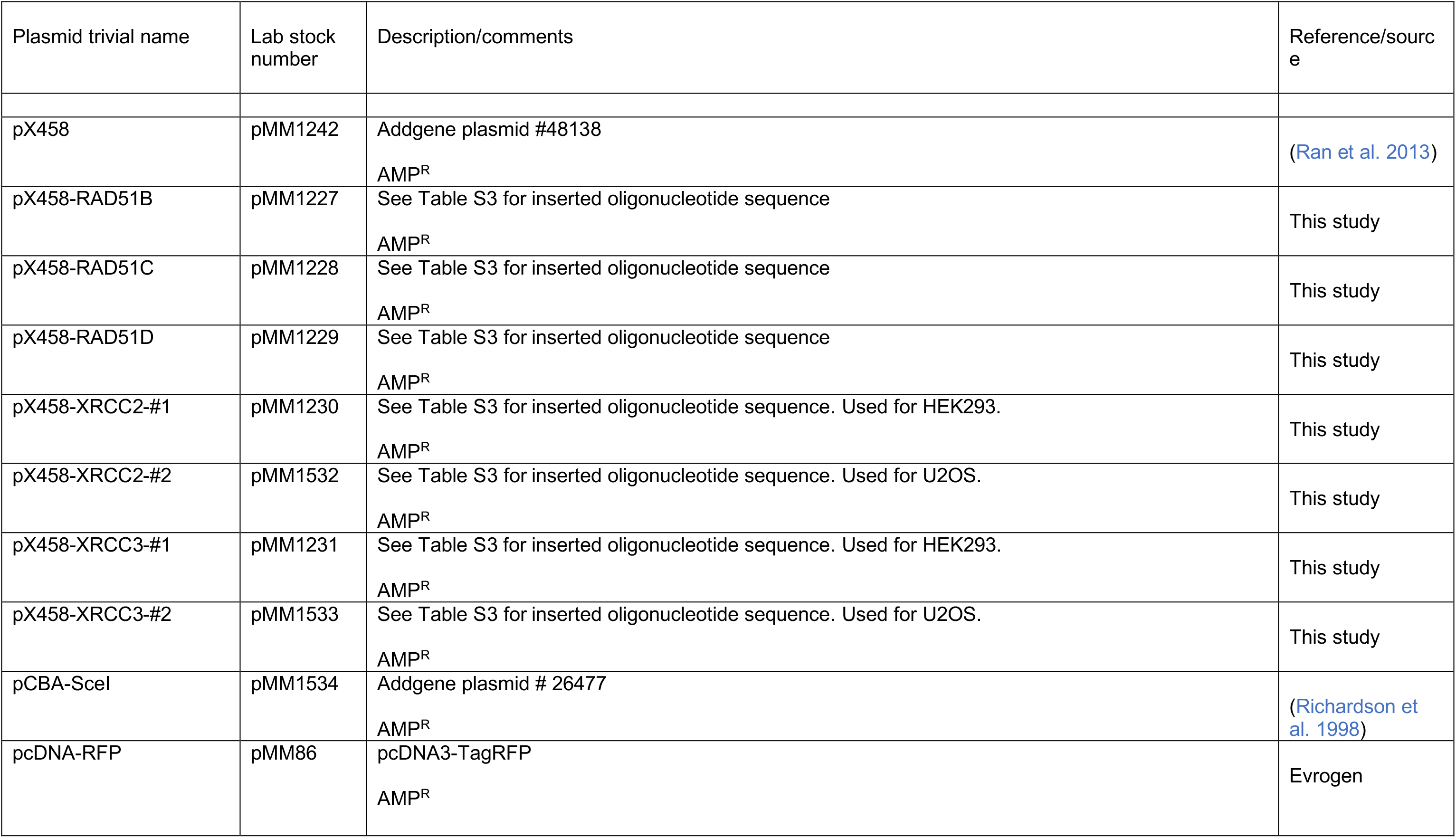
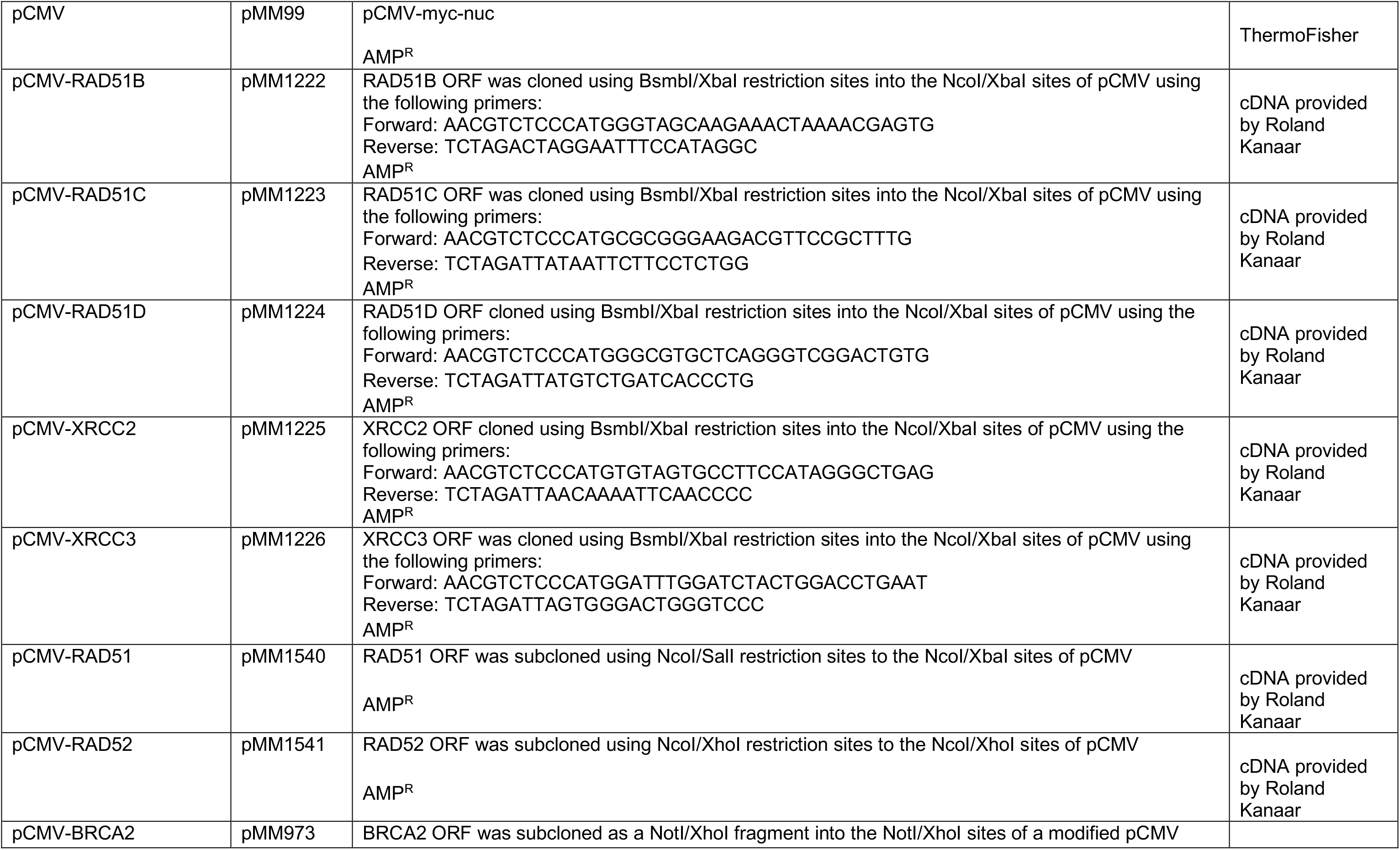
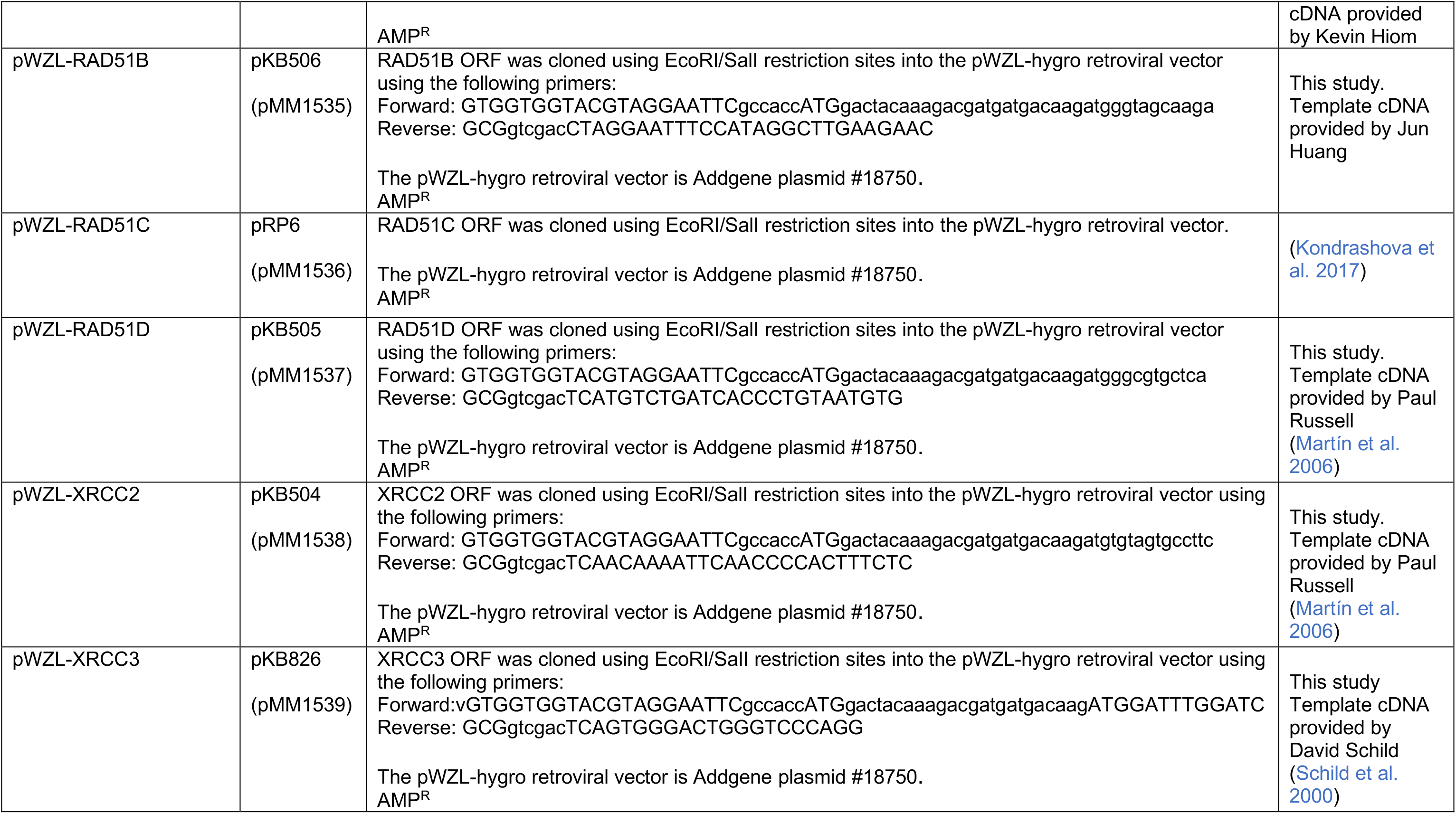
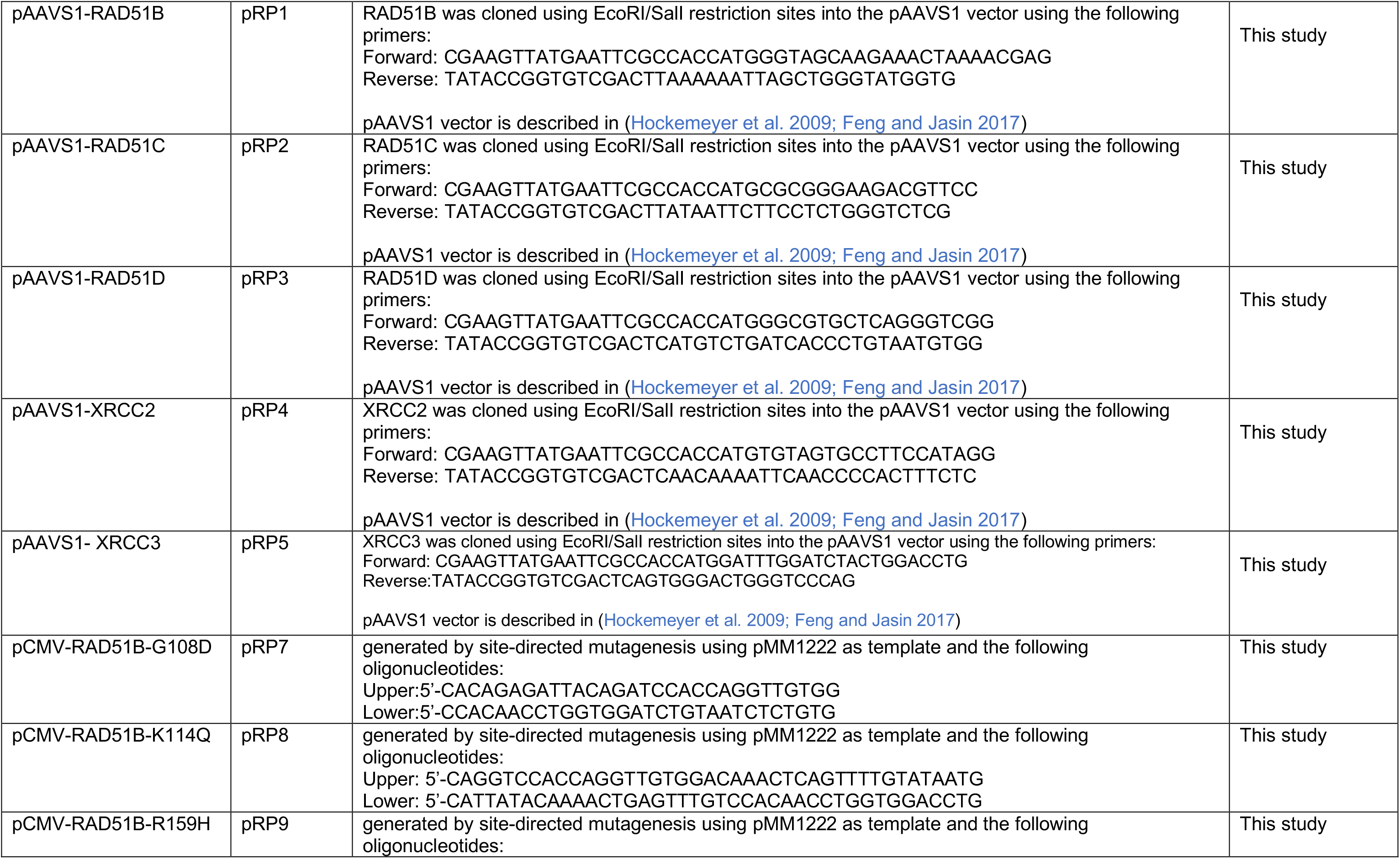
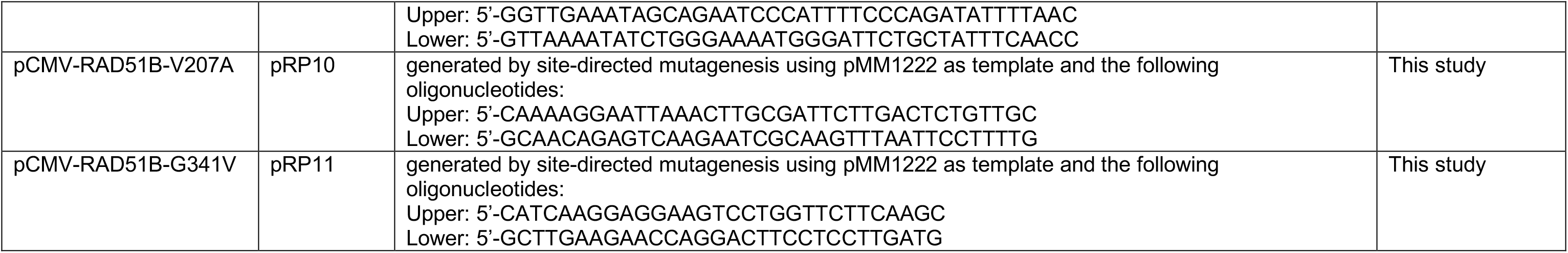
Plasmids used in this study.

**Figure S1.**
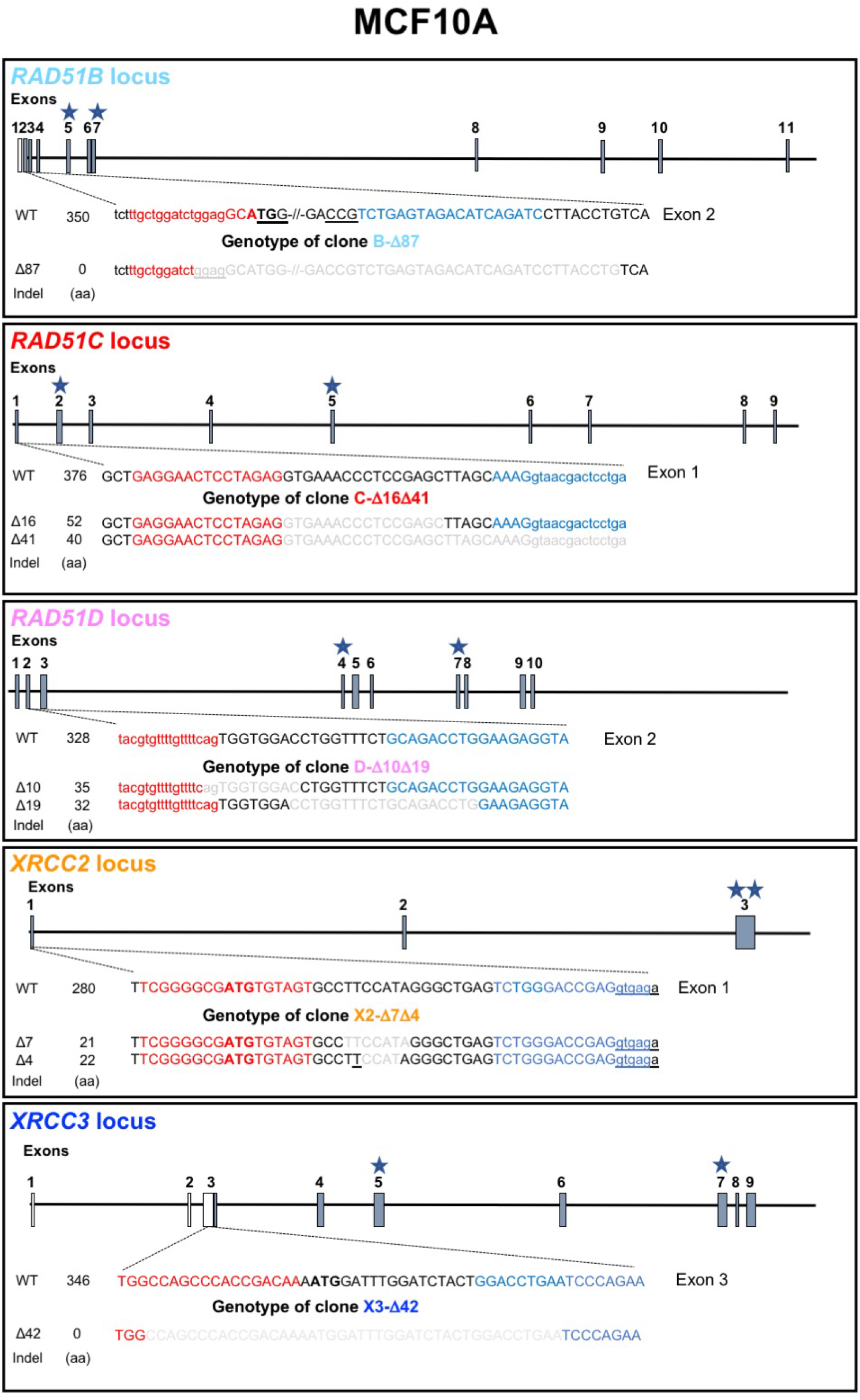
Related to Figure 1C. Inactivation of RAD51 paralogs in human MCF10A cells. Schematics showing the genomic locus for each RAD51 paralog where filled and clear exons represent coding and non-coding exons respectively (see Table S1 for reference to the Ensembl transcript). Stars indicate the locations of the sequences encoding the Walker A and B domains. DNA sequence in red and blue indicates the left and right gRNA binding sites and the PAM (underlined) for RAD51B and left and right TALEN recognition sequences for RAD51C, RAD51D, XRCC2 and XRCC3. ATG is shown in bold. Underneath these are the genotypes of the mutant cell lines used in this study with DNA sequences in gray highlighting the indels.

**Figure S2.**
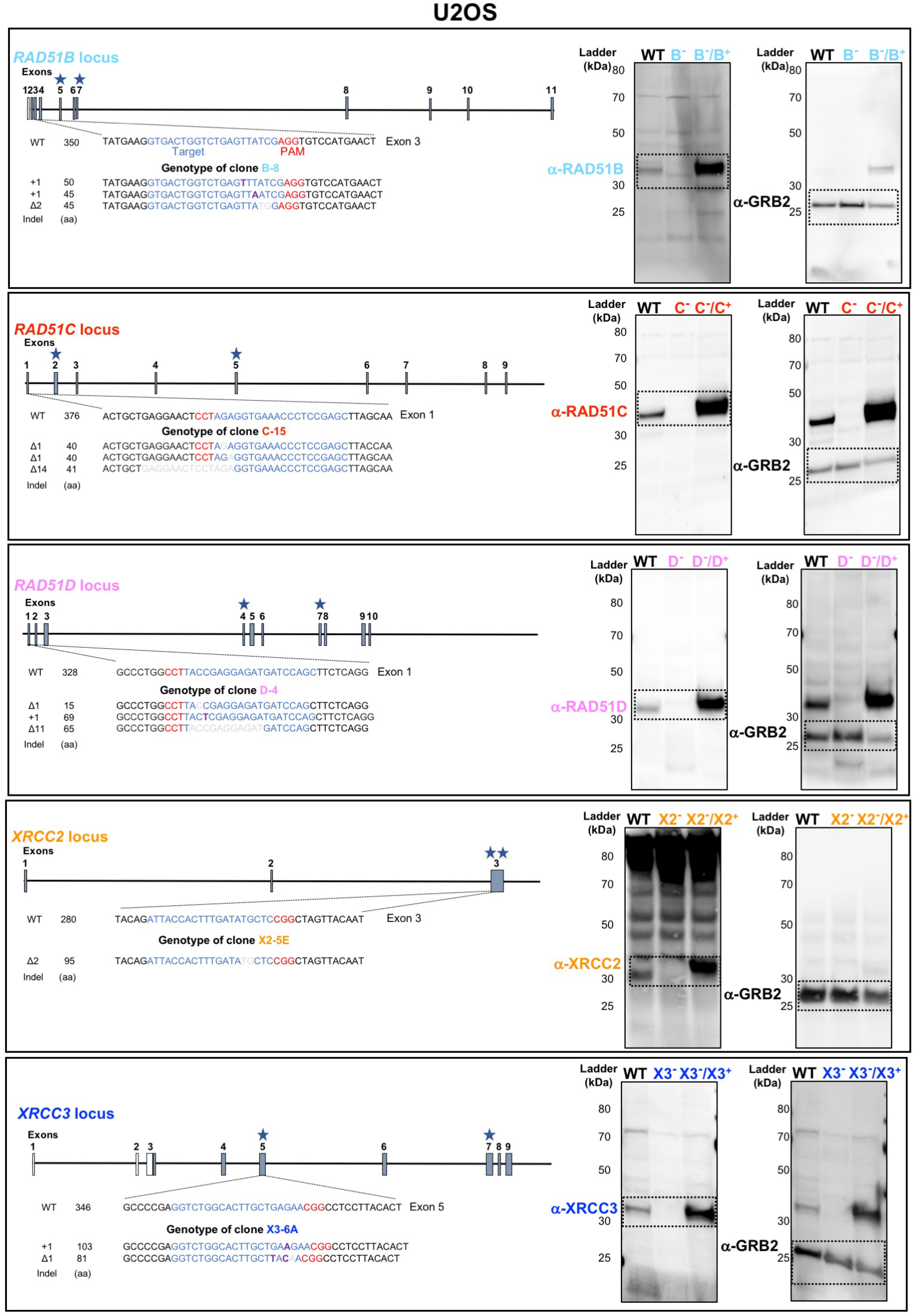
Related to Figure 1D. Individual disruption of the classical RAD51 paralogs in U2OS cells. For each RAD51 paralog, a panel is presented to show the organization of the genomic locus (see Table S1 for reference to the Ensembl transcript) and the location of the targeted site by the gRNA (blue) and the PAM (red) early in the coding sequences used for CRISPR-Cas9 genome editing (top). Stars indicate the locations of the sequences encoding the Walker A and B domains. The genotype of the mutant cell line with the indels and the size of the predicted truncated polypeptide (a.a.) are shown below. Full-size immunoblots of crude cellular extracts from wild-type cells, mutant cells and mutant cells stably complemented with a retroviral construct expressing the corresponding wild-type allele are shown on the bottom right side. GRB2 was used as loading control. See Table S1 for clone names.

**Figure S3.**
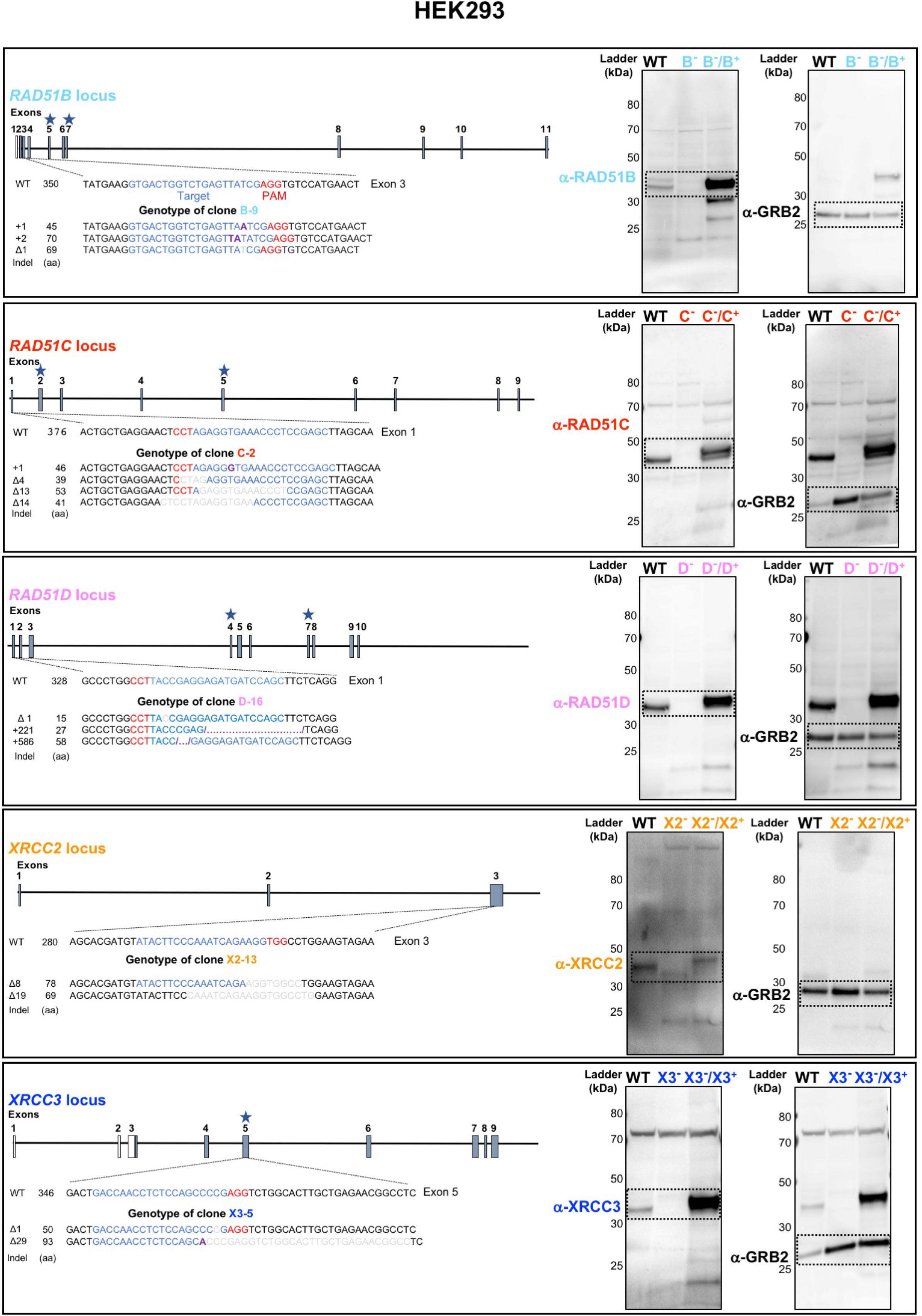
Related to Figure 1E. Individual disruption of the classical RAD51 paralogs in HEK293 cells. The legend is the same as for Figure S2.

**Figure S4.**
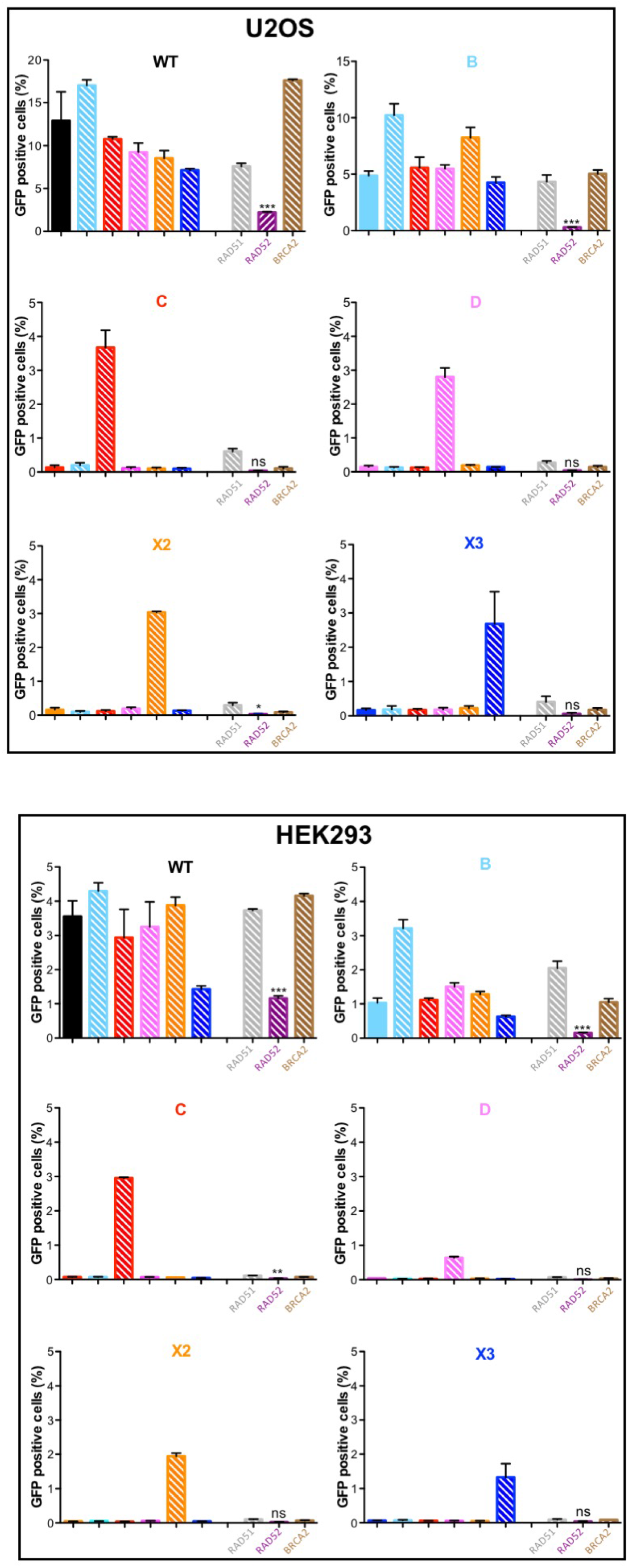
Related to Figure 3. RAD51 paralog disruption lead to homologous recombination deficiency. U2OS and HEK293 wild-type and RAD51 paralog mutant cells were transfected with I-SceI-expressing plasmid and with plasmids expressing RAD51 paralogs, RAD51, RAD52 or BRCA2 cDNAs under the control of the strong cytomegalovirus promoter. The frequencies of GFP positive cells were measured 72 h post-transfection. Data are presented as means +/− SD from at least three independent experiments. Effect of RAD52 expression was analyzed by performing unpaired one-way ANOVA followed by Tukey’s tests between results from cells transfected or not with the RAD52 expression plasmid. ns p > 0.05, * p ≤ 0.05, ** p ≤ 0.01, *** p:s; 0.001.

**Figure S5.**
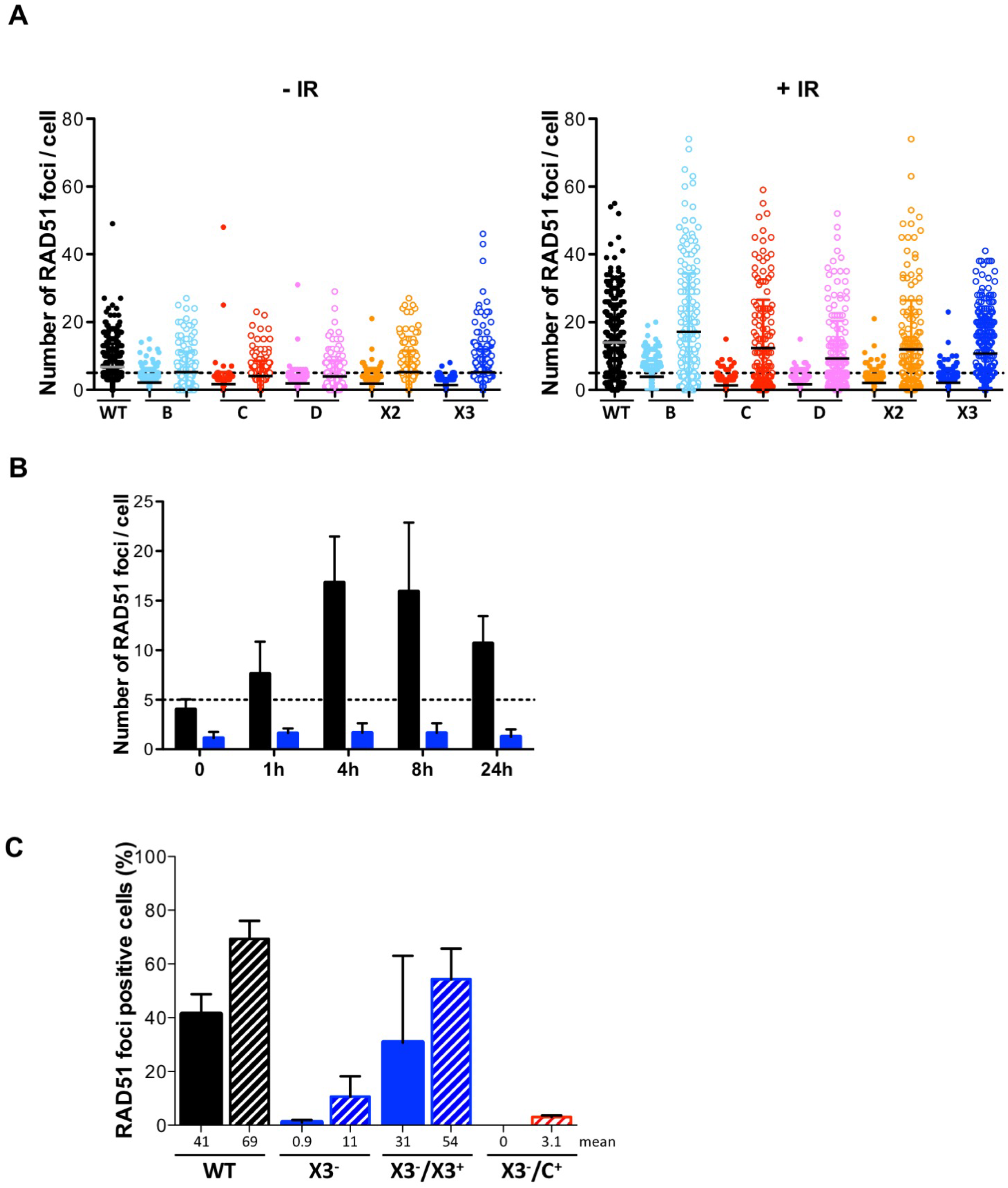
Related to Figure 4. RAD51 focus formation is defective in RAD51 paralog disrupted U2OS cells. **(A)** Quantification of RAD51 nuclear focus formation by immunofluorescence in wild-type, RAD51 paralog disrupted and stably complemented RAD51 paralog disrupted U2OS cells exposed to 0 or 4 Gy and after 4 h of recovery. Pool of all the data collected from three experiments for each cell line and condition is presented in the dot plot. Sample sizes from left to right are n= 323, 293, 186, 269, 179, 277, 173, 275, 167, 277, 279 for (-IR) condition; and n= 280, 260, 190, 276, 162, 290, 181, 261, 170, 299, 290 for (+IR) condition. **(B)** Time course of RAD51 nuclear focus formation by immunofluorescence in wild-type (black) and XRCC3 mutant (blue) U2OS cells exposed to 4 Gy and incubated up to 24 h. At least 100 nuclei were scored for each time point. Results are presented as means +/− SD from three independent experiments. **(C)** Quantification of RAD51 nuclear focus formation by immunofluorescence in wild-type, *XRCC3* mutant, and *XRCC3* mutant U2OS cells stably complemented with *XRCC3* or *RAD51C* cDNAs, respectively. Cells were exposed to 0 (filled bars) or 4 Gy (hatched bars) and scored after 4 h recovery. Data for WT, X3 and X3-/X3+ are reported from Figure 4C for comparison to the X3-/C+ experimental samples. In the latter case, two experiments were performed scoring at least 50 nuclei per experiment where in total, 125 and 132 images were analyzed for 0 and 4 Gy conditions, respectively. The data are presented as means +/− SD from the two experiments.

**Figure S6.**
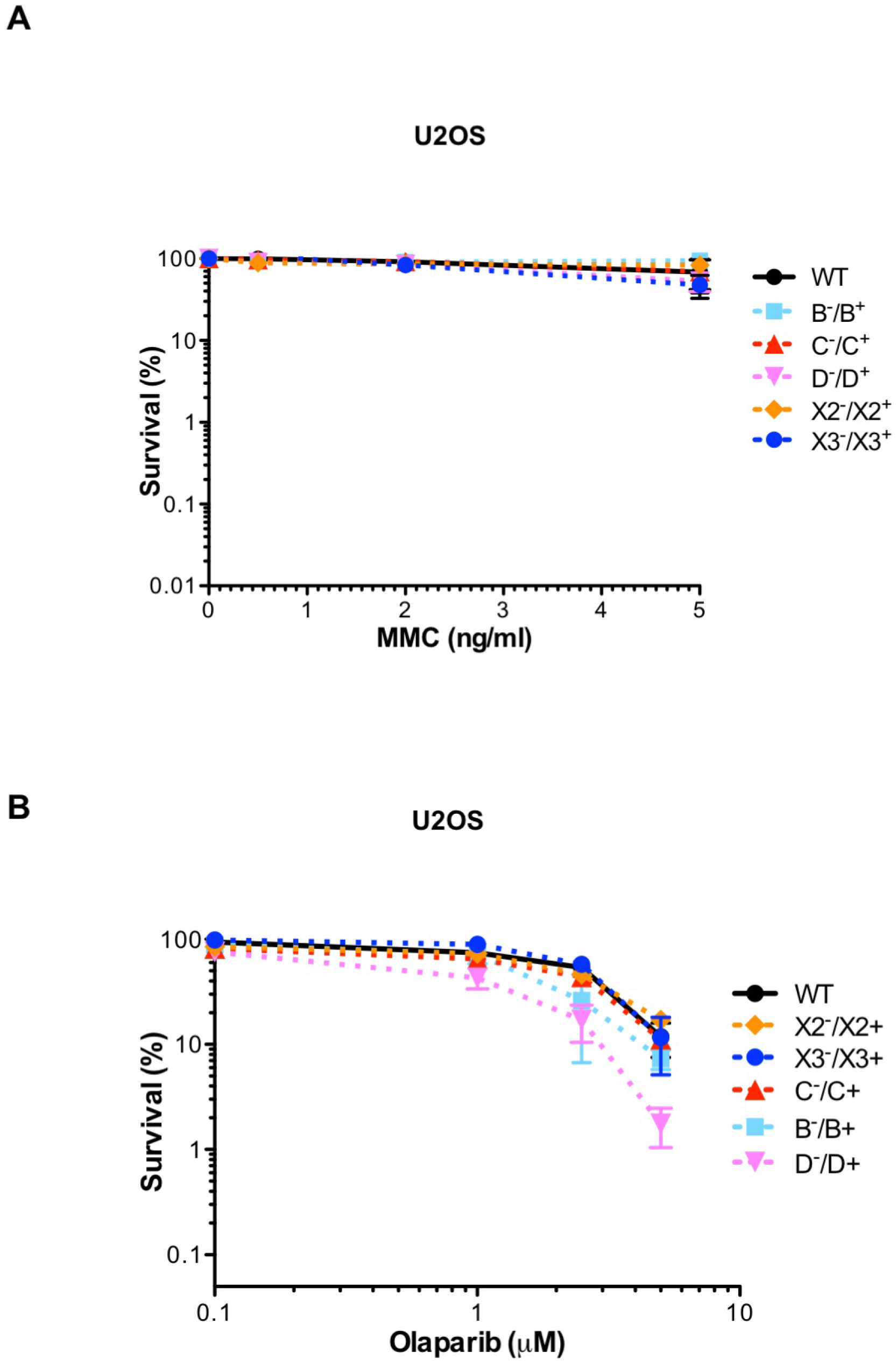
Related to Figure 5. RAD51 paralog disruption sensitizes U2OS cells to mitomycin C and olaparib. Survival curves obtained by clonogenic cell survival assays after treatment of exponentially growing U2OS cells with indicated doses of **(A)** mitomycin C (MMC) or **(B)** olaparib. Analyses of the mutant cells stably complemented with a retroviral construct expressing the corresponding wild-type allele are shown. Results are presented as means +/− SD from at least three independent experiments.

**Figure S7.**
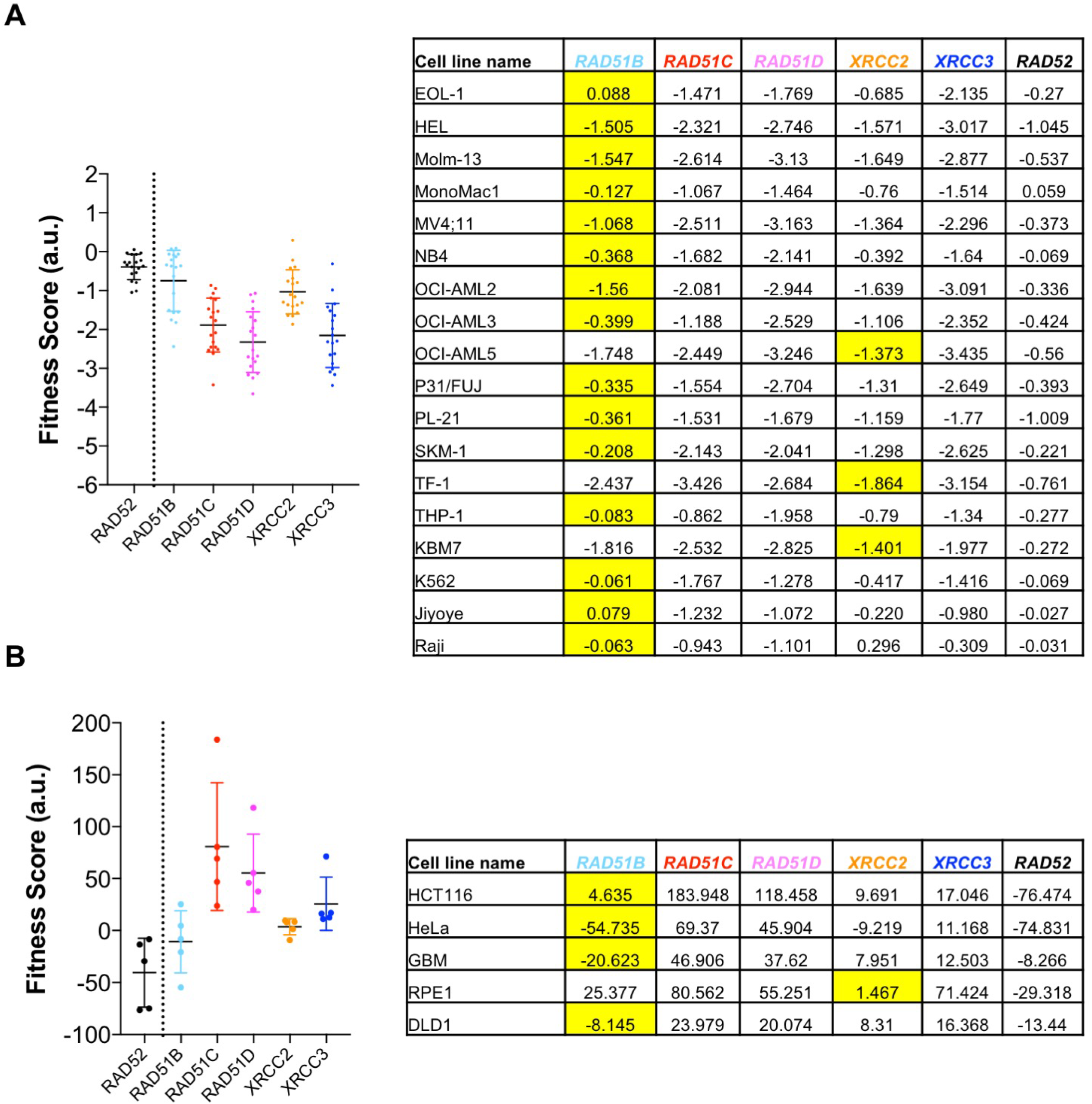
Related to the Discussion. Growth fitness of human cell lines after RAD51 paralog CRISPR-Cas9 targeting. Comparison of fitness scores expressed in arbitrary units from 18 human cell lines **(A)** and 5 human cell lines **(B)** after *RAD52* and RAD51 paralog CRISPR-Cas9 targeting predicts that disruption of *RAD51B* is more similar to *RAD52* disruption than to the disruption of the other RAD51 paralogs in terms of cell survival. Raw CRISPR-Cas9 scores after *RAD52* and RAD51 paralogs targeting from various genetic screens are shown on the right of each panel. Yellow highlights indicate the highest relative fitness score.

**Figure S8.**
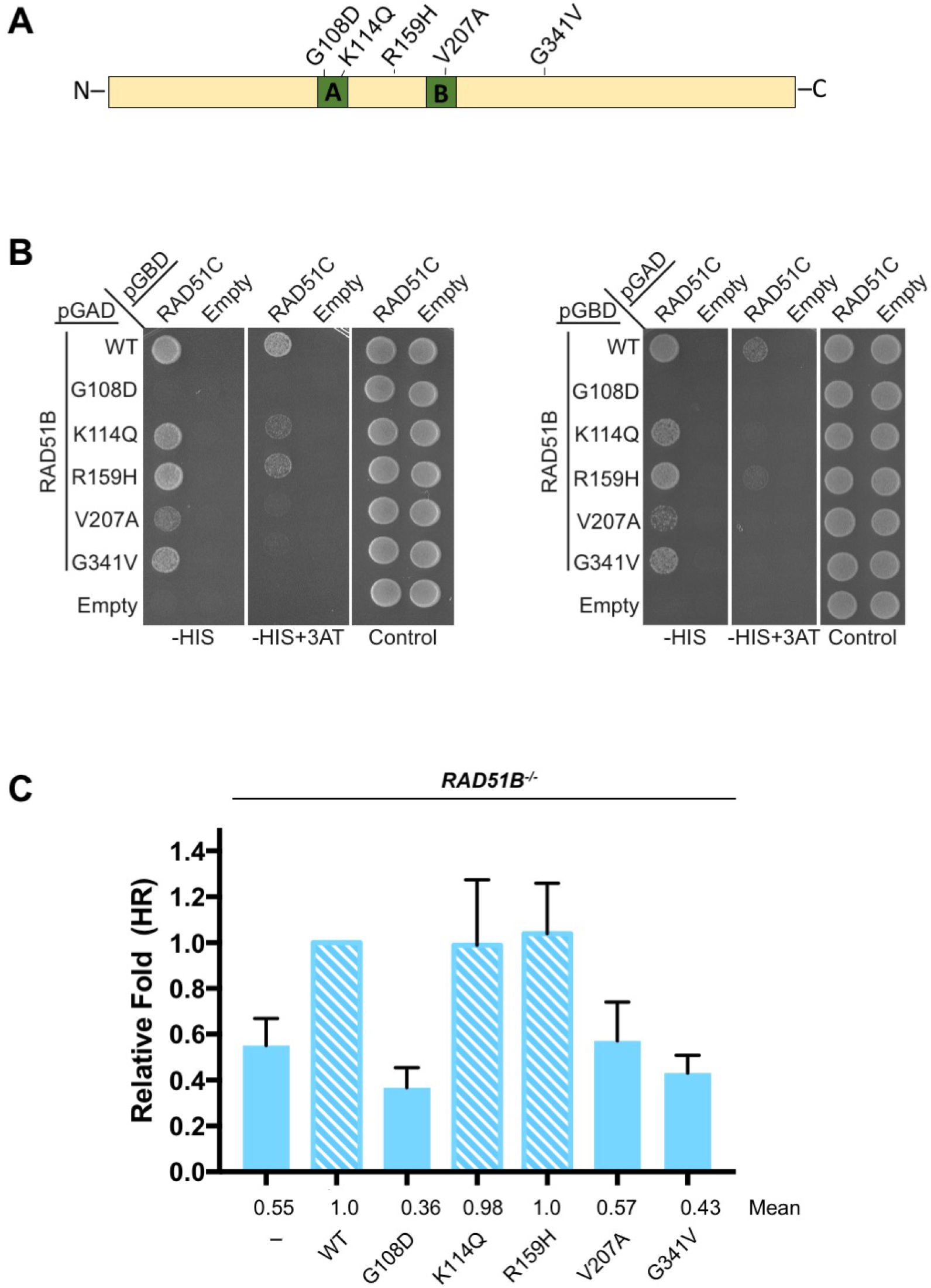
Related to the Discussion. Walker A and B domains of RAD51B are essential for its function in HR. **(A)** Schematic showing RAD51B with cancer-associated mutations analyzed in this study. **(B)** Identification of RAD51B point mutants that disrupt yeast 2-hybrid (Y2H) interaction with RAD51C. RAD51B and the corresponding point mutants were expressed in Y2H plasmids containing the GAL4 activating domain (AD) or binding domain (BD) and co-transformed with plasmids expressing RAD51C and the alternative GAL4 domain, as indicated. Empty AD and BD vectors were used as negative controls. Growth on SC-Leu-Trp-His (-HIS; less stringent) or SC-Leu-Trp-His+3AT (-HIS+3AT; more stringent) indicates a Y2H interaction whereas growth on SC-Leu-Trp (Control) indicates equal cell loading. **(C)** *RAD51B^-/-^* HEK293 cells were transfected with I-SceI-expressing plasmid and with plasmid expressing RAD51B wild-type or the mutant protein. The frequencies of GFP positive cells were measured 48 h post-transfection. The data is represented as the fold increase. Data are presented as means +/− SD from five independent experiments.

